# Ultraslow entorhinal oscillations shape spatial memory through grid cell drifting

**DOI:** 10.64898/2026.03.13.711323

**Authors:** Luca Sarramone, Matías Presso, José A. Fernandez-Leon

**Affiliations:** NeuroAI Lab, Fac. Cs. Exactas-INTIA, Universidad Nacional del Centro de la Provincia de Buenos Aires (UNCPBA), Tandil, Buenos Aires, Argentina; Consejo Nacional de Investigaciones Científicas y Técnicas (CONICET), Buenos Aires, Argentina; CIFICEN (CONICET–CICPBA-UNCPBA), CCT-Tandil, Buenos Aires, Argentina; Comisión de Investigaciones Científicas de la Provincia de Buenos Aires (CIC), Buenos Aires, Argentina

**Author notes:** Corresponding authors (main author); (senior author); NeuroAI Lab: https://neuro-ai-lab.intia.exa.unicen.edu.ar/.

**Keywords:** grid cells, spatial navigation, path integration, place cells, memory

## Abstract

**Context:** Grid cells in the medial entorhinal cortex (MEC) of head-fixed mice exhibit ultraslow (<0.01 Hz) oscillations (USO) during walking in a 1D running wheel in darkness. It was proposed that these oscillations may have a connection with navigational behavior.

**Problem:** There is no clear link between the functional role of these oscillations and path integration, a fundamental navigation strategy used by animals to calculate their current position and orientation by continuously summing self-motion cues.

**Hypothesis:** Given the synaptic projections from MEC to the hippocampus, we hypothesized that ultraslow oscillations have a role in linking spatiotemporal memories acquired during navigation.

**Methodology:** A realistic computational model of entorhinal-grid with ultraslow oscillations and hippocampal-place cells is proposed using synaptic plasticity between cell types, sustaining path integration of a rodent-like simulated animal.

**Results:** Ultraslow oscillations induced persistent changes in the grid cell dynamics, represented as a positional drift of grid fields. Such drift resulted in position estimation errors but generated new grid-place cell associations when combined with synaptic plasticity.

**>Discussions:** Ultraslow entorhinal oscillations were found to shape spatial memory through grid cell drifting, which could serve as a mechanism for flexibly accessing different spatial memories during navigation.

**HIGHLIGHTS:**

- Path integration dynamics hide ultraslow oscillations despite coexistence.
- Ultraslow oscillations significantly degrade position estimation in path integration.
- Grid and place fields drift after the effect of ultraslow oscillations.
- New spatial memories were created as a result of the ultraslow oscillation drift.
- Ultraslow oscillations enable dynamic access of different spatial memories

## INTRODUCTION

Animals navigate through the environment using an internal navigation system localized primarily in the hippocampus and medial entorhinal cortex (MEC) ^1–4^. Grid cells in the MEC are known for firing periodically as an animal navigates in space, creating a triangular pattern that tiles the environment and is thought to support position estimation ^5,6^. Together with hippocampal place cells, they represent the foundational components of the “cognitive map”, a mental representation of space used by animals to navigate ^7,8^. Grid cells contribute by constantly updating this map through path integration (PI, **Fig. 1B**), the ability to estimate the current position based on a known starting point and proprioceptive cues encoded as a velocity signal ^9–12^. Meanwhile, place cells’ contribution is to reduce the estimation error of path integration using memories of previously visited locations, anchored to environmental cues ^13–15^.

**Figure 1.**
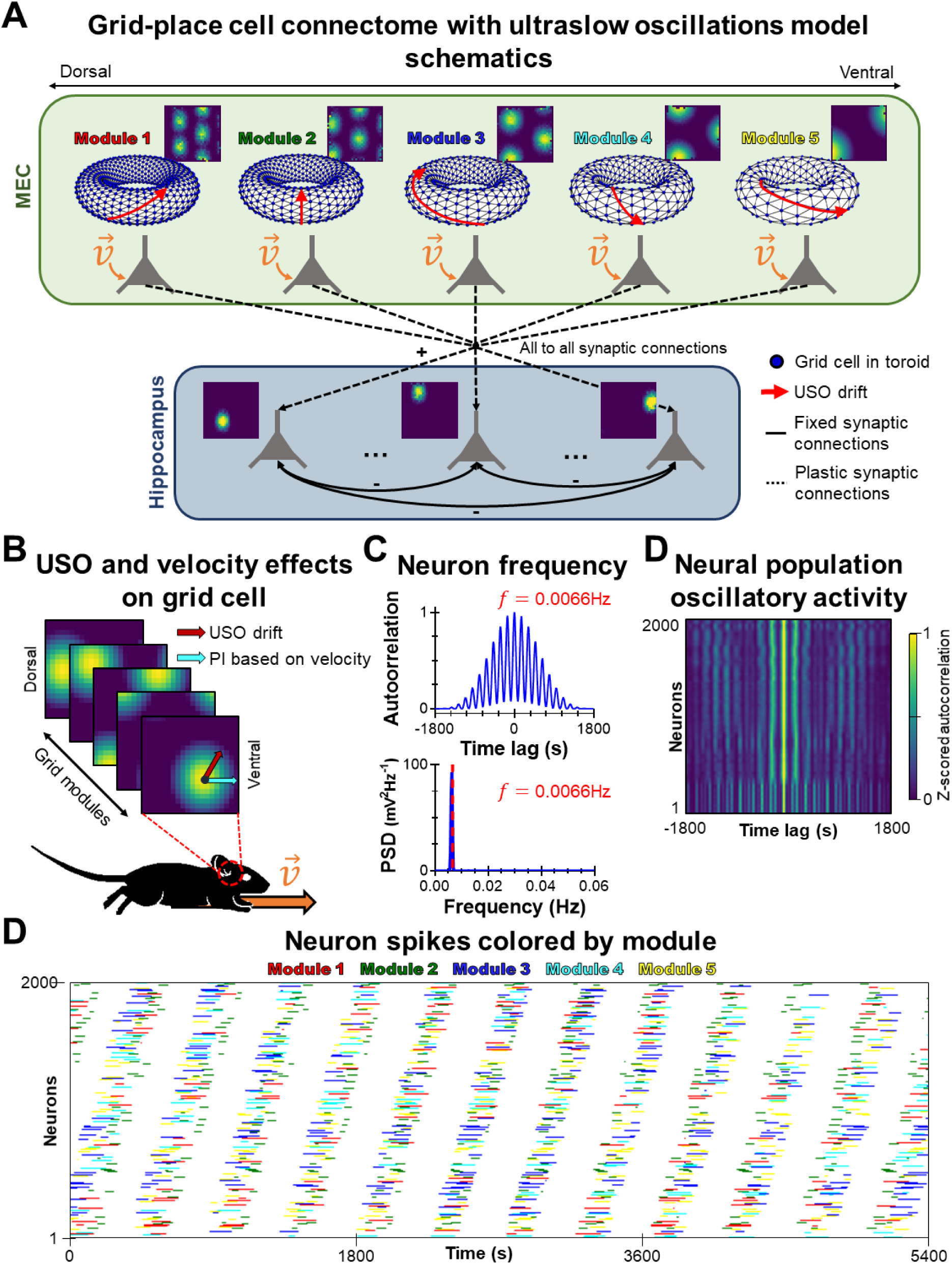
Experimental set-up and initial results. **A.** Schematics of the entorhinal grid cells and hippocampal place cells model. Five grid cell modules were used, each organized as a torus, with scales that increment from dorsal to ventral modules. All grid cell modules receive the same velocity signal (orange arrow). All grid cells from each module were interconnected with local excitatory and distant inhibitory connections. No connections between modules were established, so grid module coordination is achieved by the shared velocity input. In each module, a different synaptic weight path (red arrow in torus) was imprinted, responsible for the ultraslow oscillation (USO) effect. Place cell activity depended solely on grid cell activity. Synaptic weights connecting grid cells with place cells were trained using Hebbian learning (dashed black arrow). Place cells are interconnected with fixed inhibitory synaptic weights. **B.** Schematics of the forces acting on the grid cell dynamics. As the simulated animal moves, the activity of the grid cells is updated by the velocity (orange arrow) of the animal via path integration (PI, cyan arrow) and by the path imprinted in the synaptic weights that causes the ultraslow oscillation (USO, red arrow). The directions of both forces may or may not match in direction. **C.** Example of autocorrelation (top) and PSD (bottom) calculated over the activity of one neuron. The dotted red line signals the frequency of the oscillation. **D.** Stacked z-score correlation of the cell activity from all the neurons in the network (2000 neurons total), organized by maximum PSD activity. The pattern shows that all neurons fire at a similar frequency. **D.** Color coded raster plot of grid cell population activity over time, sorted by the PCA criterion (see methods). The simulation lasted 30 minutes (1800 s), and a total of 2000 neurons were registered. The frequency of the oscillation matched the frequency shared by the neurons in C. Grid cell from the same modules were represented by the same colors.

Grid cells in the MEC exhibit a minute-scale oscillatory pattern (<0.01 Hz) when mice run on a wheel in darkness with reduced sensorial input ^16^. These oscillations were intrinsic to the grid cell network, as inputs from place cells and other sensory-driven sources should be largely suppressed. Importantly, there is no evidence yet of the frequency of these oscillations matching the time scale of path integration, a mechanism related to grid cells ^17,18^. However, grid cells are also associated with memory processes ^15,19,20^, and a similar ultraslow phenomenon has been previously reported to affect the hippocampus ^21^, a brain area strongly associated with spatial and episodic memories. Synaptic projections from the MEC to the hippocampus have been previously reported ^22–25^, hinting that the oscillatory process observed in the hippocampus could originate in the MEC. Based on this evidence, we hypothesize that MEC ultraslow oscillations (USO) influence hippocampal place cells and have an effect on spatial memories rather than supporting path integration. Functional relations between grid and place cells during navigation have been extensively discussed, with grid cells providing support when sensorial input is absent, and place cells providing a mechanism for error resetting ^13,26,27^.

Based on our previous work ^13,28^, we developed a grid cell continuous attractor model (CAN) that exhibited ultraslow oscillations. Grid cell CAN models are based on the assumption that grid cell activity lies on a toroidal manifold supported by structured recurrent connectivity ^29,30^. When the synaptic connections are perfectly balanced, the network remains in a stable state unless it is perturbed by an outside force (e.g., a speed signal) ^31,32^. However, an internal driving force emerged when alterations to the network synaptic weights were induced, generating an oscillatory signal in the grid cell dynamics independent of external input. The model was capable of path integration using an external velocity signal. This input could be suppressed to isolate the oscillatory dynamics and study their effect on the navigation behavior. In addition to the grid cell model, a place cell network was trained using Hebbian plasticity and the grid cell activity recovered during navigation. As the place cell population activity was solely determined by the grid cell activity, we used it to study the effects of ultraslow oscillations in spatial memory.

Our work serves as a theoretical proof of how ultraslow oscillations can emerge from a simple synaptic modification mechanism over a CAN model. We extensively discuss the role of ultraslow oscillations in path integration and spatial memory recall during navigation, providing a foundation for future investigations.

## RESULTS

### Synaptic weights tuning is sufficient to generate an ultraslow oscillation

We created a virtual environment in which a simulated rodent-like animal moved while updating a continuous attractor network for position estimation based on grid cells ^28^. The grid network was composed of five grid modules, each with *N* = 400 grid cells. The scale of grid cells was the same among neurons from the same module, but increased linearly between modules by a gain factor of 0.02 (see Methods: *Grid scale and gain relation*), mimicking the dorso-ventral organization reported in the MEC ^33^. The network generated a single packet of activity (or “bump”) that traveled on a toroidal manifold by integrating the simulated animal’s velocity through path integration. Once the animal stopped moving, so would the activity bump, and the grid cell population activity remained fixed. At the same time, a place cell network was trained using Hebbian learning and the grid cell activity during exploration. These place cells worked as an indicator of spatial memories ^8,15,34^, and their place fields only emerged after the animal explored the whole environment (**Fig. 1A**).

To implement the reported ultraslow oscillation ^16^, we modified the attractor network by adding a directed force to the activity bump. This force was implemented as a small, structured perturbation to the synaptic weights. While the original network produced a single stable attractor, applying an identical perturbation to all synaptic connections created a gradient that induced a slow drift around the torus (**Fig. 1A**; see Methods). This perturbation was parametrized by a direction (*θ*) and a strength of the alteration in the synaptic weights (*ρ*) (see Methods), which determined the direction and speed of the activity bump drift (**Fig. 1B**). The oscillation was solely affected by the direction and strength of the synaptic weight gradient, and not by other parameters such as the module scale or initial state of the grid network. Each module had its own set of parameters, which could be the same or different from those of other modules. Both the velocity signal and the ultraslow oscillation could coexist, allowing the activity bump to drift even after movement ceased.

We assessed whether the oscillatory behavior of our model resembled MEC oscillations by running a 30-minute simulation at a sampling rate of 7.73 Hz (13914 samples) as in ^16^. During this period, the velocity signal was inhibited to isolate the drift effect from the path integration dynamic (see Methods). An initial analysis of one neuron of each module showed that the autocorrelated activity of each neuron had multiple side peaks, indicating oscillatory activity, also confirmed by applying a power spectral density (PSD) analysis (**Fig. 1C**, **Suppl. Fig. S1A**). We tested if all neurons from the same module shared similar oscillatory behaviors, as they have the same *θ* and *ρ* parameters. By computing and stacking the correlated z-score by module, we confirmed that all neurons from the same module shared comparable oscillatory behaviors, with frequencies near 0.0066 Hz (**Suppl. Fig. S1B**). To further confirm the oscillatory behavior per module, we projected the activity of each module into a 2D-plane using PCA and observed a cyclical pattern (**Suppl. Fig. S1C**), similar to the one reported in ^16^.

To reproduce the population-wide synchrony reported by Gonzalo-Cogno and colleagues^16^, we systematically explored combinations of the *θ* and *ρ* parameters from all modules and grouped them by their resulting oscillation frequencies (**Suppl. Fig. S2A**). Larger *ρ* values yielded faster drift, whereas *θ* values farther from multiples of 90° generated longer toroidal paths and thus slower oscillations. Interestingly, despite reaching high *ρ* values during the parameter exploration, the generated frequencies tended to group in the lower range, with the highest frequency being ∼0.3 Hz (**Suppl. Fig. S2A**). This result suggested that oscillations generated using weight tuning were inherently slow, with frequencies typically ranging between 0.0002 Hz and 0.123 Hz, with some occasionally reaching higher values (<0.3 Hz). We then selected five parameter sets from the 0.0066 Hz frequency group and verified that the full population was synchronized using the z-score autocorrelation (**Fig. 1D**). This frequency was chosen because it matched the frequency reported ^16^, but other frequencies could be used. The final parameters chosen to replicate the 0.0066 Hz frequency per module were *ρ* = [0.2, 0.45, 0.2, 0.2, 0.3] and *θ* = [290°, 140°, 190°, 240°, 210°]. These parameters were the ones used in the rest of the experiments described in this paper, unless stated otherwise.

We then tested whether the oscillatory population activity formed the characteristic ultraslow pattern. Neurons were ordered using two criteria (see Methods). The first method consisted of cross-correlating neural activity within small time lag windows to identify near co-activations of neurons (**Suppl. Fig. S2B**). The second method was based on principal component analysis (PCA), and projected the neural activity into a new *R*^2^ space where a single unifying criterion was used to organize the firing of all cells (**Suppl. Fig. S2C**). Both methods produced a clear ultraslow oscillation pattern, but the PCA method was much more computationally efficient, so it was chosen as the default method for the rest of the analyses in this work. Notably, neurons from the same grid module did not fire at consecutive phases of the oscillation (**Fig. 1D**). Considering that the modeled grid cell modules were organized following the scale increments observed in the dorsal-ventral axis of the MEC ^33^, results indicated that the oscillation did not correspond to a traveling wave, consistent with in vivo results ^16^. In summary, small, structured perturbations to the synaptic weights of a continuous attractor network were sufficient to generate ultraslow oscillations based on a continuous attractor network.

### Path integration suppresses ultraslow oscillations at the cost of reduced accuracy

We investigated the circumstances under which ultraslow oscillations were evidenced. Four experimental conditions were simulated (**Fig. 2A**): (a) head-fixed-like movement in one direction, resembling a running wheel; (b) forward and backward movement in a running wheel; (c) freely moving in a 2D arena enclosed by walls; and (d) open-field free movement. Each condition imposed different constraints on the simulated animal’s movement. While the first condition replicated the experimental setup of ^16^, with mice’s movements restricted to forward movement or immobility, the remaining conditions progressively increased the degrees of freedom, from bidirectional 1D motion to unrestricted 2D exploration. For each condition, 30 different runs were tested to reduce the effects of randomness during exploration. Each run consisted of three stages: 30 minutes of immobility, 30 minutes of movement, and a final 30 minutes of immobility. This design allowed us to examine the ultraslow oscillation effect before, during, and after the path integration process. During immobility, the only force affecting the grid cells dynamics was the ultraslow oscillation, making it more detectable. Importantly, stopping movement rather than inhibiting the speed signal prevented the accumulation of position estimation errors due to path integration stopping while the animal kept moving. This error accumulation would otherwise confound the analysis. However, both the path integration process and the ultraslow oscillation could be active simultaneously.

**Figure 2.**
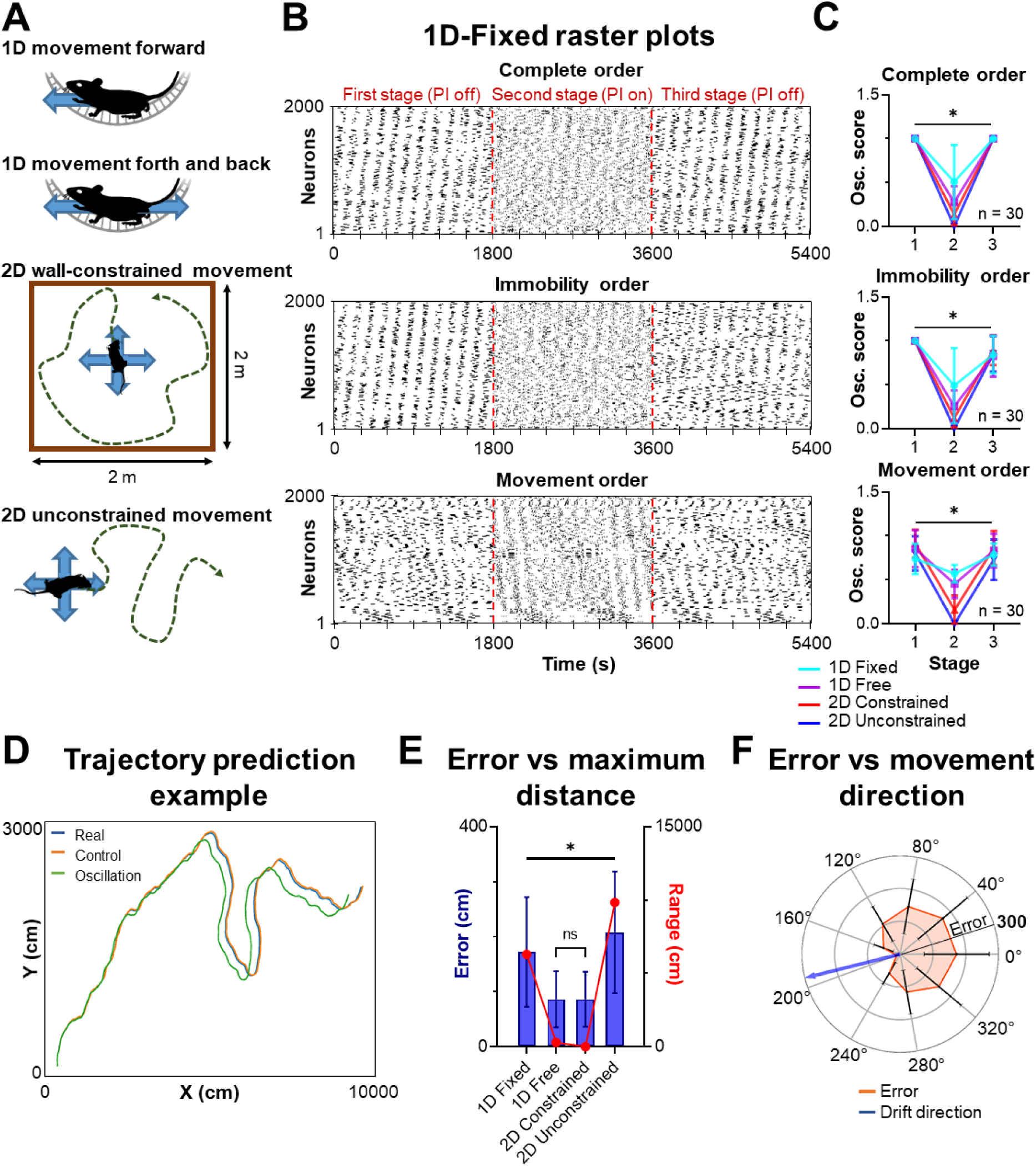
Ultraslow oscillations increase error accumulation during path integration. **A.** Schematics of the four experimental conditions tested. From top to bottom, the movement possibilities for each simulated animal are: (I) Move forward or stop; (II) Move forward, backwards, or stop; (III) Move in any direction within a 2m^2^ arena; and (IV) Free, unconstrained movement in any direction. **B.** Raster plot example of neuron activity over time, sorted by three different criteria. From top to bottom: (I) Based on the neuron activity through the whole experiment; (II) Based on the neuron activity of only the first third of the experiment; (III) Based on the neuron activity of only the second third of the experiment. **C.** Oscillation score comparison of each stage of the experiments. Significant differences were found in all and across all criterions (Dunn’s multiple test, *p* < 0.001, see Suppl. Table). **D.** Example trajectory during 2D unconstrained exploration. The same trajectory was estimated with (green) and without (orange) ultraslow oscillations, and compared to the real trajectory (blue). **E.** Mean estimation error (blue, left y-axis) across experiments under oscillation conditions compared to the maximum distance reached at the end of each experiment (red, right y-axis). Significant differences were observed among all experiments (One-way ANOVA, *p* < 0.001), except between the 1D free and 2D constrained conditions, which did not differ significantly (Tukey’s multiple test, *p* > 0.05). **F.** Polar histogram showing the mean positional estimation error for different trajectories’ direction when an oscillation drift with a phase of 194° was present. The radius indicates estimation error in cm and is divided into sections of 100 cm delimited by grey rings. The blue arrow indicates the drift direction, and the black lines at each point show the standard deviation. Significant differences were found among all directions (One-way ANOVA, *p* < 0.001) and between all directions and the one with the lowest error (Dunnett’s multiple test, *p* < 0.001).

We applied frequency analyses over an ordered neural population signal created by computing the sine of the oscillation phase at each time step (see Methods). The spectrograms calculated over this signal revealed a clear oscillatory component at the ultraslow frequency (0.0066 Hz) during stillness across all conditions. However, during active movement when path integration took place, the ultraslow component decreased in power or vanished entirely (**Suppl. Fig. S3A**). Comparing the first (**Suppl. Fig. S4, first column**) and second (**Suppl. Fig. S4, second column**) stages showed that this peak markedly decreased during movement or even disappeared completely (**Suppl. Fig. S3B**). The mean power of the 0.0066 Hz frequency peak was significantly reduced during mobility compared to immobility (Wilcoxon-test, *p*<0.001, see Supplemental Table, **Suppl. Fig. S3C**).

We assessed how much of the oscillation was masked by the path integration dynamic using the oscillation score ^16^. The oscillation score depended on the order set by the PCA and the correlation order (see Methods). Three ordering criteria were used: (a) order based on the complete cell activity; (b) order based only on the first stage of the experiment, with no movement and only ultraslow oscillations; and (c) order based on the second stage of the experiment, with movement and ultraslow oscillations (**Fig. 2B, Suppl. Fig. S5**). The oscillation scores computed for each experiment indicated statistical differences among all three order criteria (Friedman test, *p*<0.001, **Suppl. Fig. S3D**), particularly between criterion (c) and the other two criteria (Dunn’s multiple test, *p*<0.01, **Suppl. Fig. S3D**). We then computed the oscillation score separately for each stage and found significant differences between them in all experimental conditions, without regard to the order criteria used (Friedman test, *p* < 0.05, **Fig. 2C**). In all three cases, the second stage was the one that exhibited the lowest oscillation score consistently. This outcome was coherent with the spectrogram results and indicated that during periods of path integration, the ultraslow oscillation dynamics were almost completely suppressed.

We then focus on the effect of the ultraslow oscillation on path integration dynamics. For each experiment, we took the neural activity corresponding to the path integration period and estimated the traveled trajectory (see Methods). We compared the grid cell activity with ultraslow oscillation (Oscillation) and without it (Control). This approach allowed us to test if, despite not being visible in the raster plots, ultraslow oscillations improved the path integration process. When the estimated trajectories were compared to the actual paths, clear deviations appeared, especially near the trajectory endpoints (**Fig. 2D**). We quantified this estimation error as the Euclidean distance between real and estimated positions for each step of the trajectory in both conditions. The accumulated error increased exponentially over time (**Suppl. Fig. S6A**), and the mean error was significantly larger in the oscillation condition when compared to the control condition (Sidak’s multiple test, *p* < 0.001, **Suppl. Fig. S6B**). This pattern held true when analyzing trajectory estimates by grid module (Sidak’s multiple test, *p* < 0.001, **Suppl. Fig. S6C**). Differences in the mean estimated error among modules in the same experiment were caused by the *ρ* parameter defined in each module. Larger *ρ* values implied larger drifts in each time step and an overall greater error accumulation for that module. Thus, adding ultraslow oscillations amplified the intrinsic drift inherent to path integration dynamics ^35,36^.

Although the increased estimated error was consistent in all experimental conditions, the magnitude of the error increase varied among them (One-way ANOVA, *p* < 0.001, **Fig. 2E**). To test which variable affected the error accumulation, we initially computed the maximum distance reached at the end and compared it with the mean estimation error (**Fig. 2E**). Results showed that the maximum distance followed the same trend as the estimation error, with 2D-unconstrained and 1D-fixed experiments producing both the greatest distances and the largest errors. In contrast, 1D-free and 2D-constrained conditions where the distance to the origin was shorter also showed much smaller errors. The same pattern was observed in the control condition, albeit with lower overall errors (**Suppl. Fig. S7A**). These results indicate that greater travel distances amplify estimation error ^37^ and that ultraslow oscillations further exacerbate this deviation.

We noted that trajectories during the oscillation stage were consistently shifted in a particular direction, prompting us to test the effect of directionality on estimation (**Fig. 2D**). To investigate whether the ultraslow oscillation pushed the position estimation along a preferred direction, independent of the animal’s movement, we ran 10 different iterations of the 1D-fixed experiment, each time rotating the direction by 40 degrees. The 1D-fixed condition was chosen because it exhibited a large estimation error but was unaffected by direction changes. We then computed the mean estimation error for both the control and oscillation conditions. While the control condition showed no particular tendencies (**Suppl. Fig. S7B**), the oscillation condition evidenced a clear minimum around the 200° angle (One-way ANOVA, *p* < 0.001, **Fig. 2F**). This result persisted when using finer directional resolutions with identical oscillation parameters (One-way ANOVA, *p* < 0.001, **Suppl. Fig. S7C**). To relate this finding to the oscillatory drift itself, we integrated the bump displacement induced by the oscillation and found a dominant drift direction of ∼194° (**Suppl. Fig. S7D**), closely matching the minimal error angle in **Fig. 2F**. Changing the directionality of the grid bump drift by adjusting the oscillation parameters produced similar outcomes (**Suppl. Fig. S7E**). Results showed that when the trajectory direction was similar to the ultraslow oscillation direction, the error was minimized.

In summary, despite ultraslow oscillations disappearing from the raster plot due to path integration taking place, their effect on the grid cell dynamics persisted. In consequence, path integration was impaired, increasing the neural drift and rapidly accumulating positional error. In this phenomenon, heading and total distance from the center were the main factors that affected the error increment.

### Ultraslow oscillations generate lasting changes in the spatial codification of grid and place cells

We tested whether the effect induced by the addition of ultraslow oscillation was reset once the oscillation input was removed. Similar to previous experiments, we simulated the animal movement for a total of 90 minutes, divided into three periods. Two identical networks were run in parallel (control and oscillation), both receiving the same velocity signal. Path integration was active during the whole experiment to capture the spatial distribution of the neurons, while the ultraslow oscillation effect was turned on only in the oscillatory network during the second stage. In all the other periods, the ultraslow oscillation was deactivated. This configuration allowed us to test whether grid cell dynamics were affected by the previous presence of ultraslow oscillations.

Results showed a rapid increase in estimation error during the second stage, consistent with the presence of an ultraslow oscillation. In all four experimental conditions, after the ultraslow oscillation was turned off, the estimation error did not reset, but its growth rate was reduced to the same level as the control network (**Suppl. Fig. S8A**), which reflected the intrinsic error of grid cells ^13^. A similar result emerged from the step-by-step correlated activity of the whole grid cell population (**Suppl. Fig. S8B**). During the first stage, the maximum correlation was achieved, but once the ultraslow oscillation was activated, the correlation fluctuated with a mean near 0, indicating that the dynamics of both networks differed. After the oscillation was turned off, correlations stabilized again but never returned to the initial values despite identical external conditions. These findings suggested a persistent change in the grid cells dynamics, induced by the ultraslow oscillations. Using the 3D PCA projection of the grid activity, we analyzed how the grid cell dynamics were altered throughout the simulation (**Fig. 3A**, **Suppl. Fig. S8C**). In all experimental conditions, the oscillatory configuration exhibited a clear difference between each stage of the experiment, with the second stage forming a transition between stages 1 and 3. In contrast, the control network dynamics remained constrained to a smaller subspace. Results indicated that despite differences in the experiments, ultraslow oscillations always generated a transition to a new grid cell dynamic.

**Figure 3.**
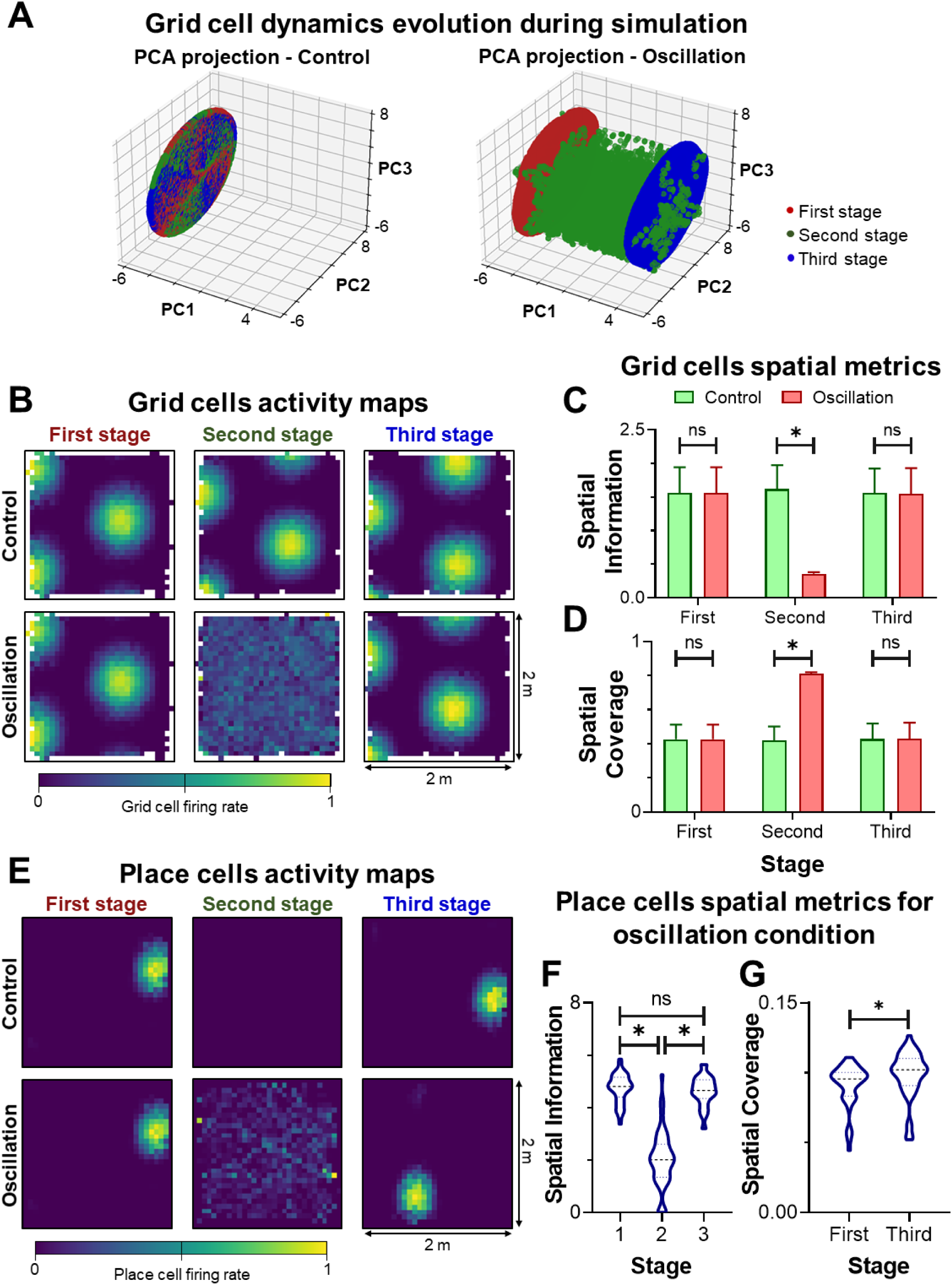
Ultraslow oscillations induce lasting changes in grid cell dynamics. **A.** Comparison of the 3D projection of grid cell activity in the PCA space for control and oscillation activity. The activity corresponds to the 1D-Fixed experiment results, and the new space where the activity is projected is the same for both conditions. Experimental stages are separated with colors: Start (no-oscillation, red); middle (oscillation, green); and end (no-oscillation, blue). **B.** Grid cell activity maps comparison for control and oscillation conditions, separated by stage. The activity corresponds to the 2D-Constrained experiment, as it is the only one where the activity map can be consistently recreated. **C.** Mean spatial information and **D.** mean effective spatial coverage of the grid cell population. Both spatial metrics were calculated by stage and experimental conditions for comparison. Significant differences in both cases were found during the second stage (middle) of the experiment (Dunn’s multiple test, *p* < 0.001). For the first (start) and third (end) stage no significant differences were found (Dunn’s multiple test, *p* > 0.001). **E.** Example of place field drift in the presence of ultraslow oscillation, divided by experiment stages. In the control condition (top row) the place field suffers a small neural drift downwards caused by path integration error. In the oscillation condition (bottom row) this drift is larger and in different directions. **F.** Place cell spatial information comparison between experiment stages. No particular alterations were found before and after the oscillation (Dunn’s multiple test, *p* > 0.05), but there were differences during the oscillatory activity stage (Dunn’s multiple test, *p* < 0.001). **G.** Effective spatial coverage of place cells before (fist stage) and after (third stage) the oscillatory period. A small increase in the spatial coverage was found between the two stages (Wilcoxon test, 0.001 < *p* < 0.01).

As long as the grid cell dynamics remained unaltered by the ultraslow oscillation, clear place fields emerged (**Fig. 1A**, **Fig. 3E first stage**), indicating the appearance of stable spatial memories. However, the observed persistent modification of the grid cell dynamics after the ultraslow oscillation period prompted us to test if new spatial representations emerge after this alteration. Place cells were defined using solely grid cell input and Hebbian learning with competition via k-WTA (see Methods). This configuration implied that any change in the grid cell spatial behavior would affect downstream place cells. We checked how the oscillations affected grid spatial behavior by generating the activity map of grid cells in the 2D-constrained condition. This condition was chosen because it reflected the hexagonal pattern of grid cells. Comparing the activity maps revealed that the grid pattern was disrupted during the second stage and re-emerged with a new phase once the ultraslow oscillation was silenced in the third stage of the experiment (**Fig. 3B**). The control network also exhibited a neural drift (**Fig. 3B, top row**), consistent with the intrinsic estimation error of grid cells ^38–40^. Importantly, this intrinsic path integration drift became visible only during path integration and when the experiment was segmented into stages; otherwise, the activity map simply showed larger and noisier grid fields (**Suppl. Fig. S9A**).

Using 2D correlations, we quantified the drift between stages 1 and 3 for both conditions. A descriptive analysis of both conditions revealed that the standard deviation of the drift in the oscillation network was larger than in the control one (**Suppl. Fig. S9B**), indicating a wider distribution of possible grid cell phases. Although the mean drift distance was not statistically different between conditions (Wilcoxon paired test, *p* > 0.05, **Suppl. Fig. S9C**), their variances diverged significantly (Pitman-Morgan test, *p* < 0.01). This result hinted that ultraslow oscillations may facilitate phase repositioning, a phenomenon observed when animals adapt to a new environment or when visual cues are diminished ^11,41^, so we measured the spatial information and spatial entropy of the grid population across the three stages of the experiment. When ultraslow oscillations were active, the spatial information was significantly reduced (Dunn’s multiple test, *p* < 0.001, **Fig. 3C**), and the spatial entropy increased accordingly (Dunn’s multiple test, *p* < 0.001, **Suppl. Fig. S9D**). This phenomenon was caused by grid cells firing in more locations of the arena during the second stage, losing location specificity but enlarging their active area. To quantify this change directly, we introduced the effective spatial coverage (ESC) metric (see Methods). After activating the ultraslow oscillation, the ESC increased dramatically, reaching arena coverages of over 80% while the control condition remained below the 50% threshold (Dunn’s multiple test, *p* < 0.001, **Fig. 3D**).

The grid cell phase shifts implied that each location in the arena was encoded differently before and after the ultraslow oscillation period. Such behavior indicated a link between the ultraslow oscillation and the encoded spatiotemporal memories during navigation. Existence of a relation between grid cell path integration and spatial memories encoded as place cells for error reduction during navigation has been previously proven ^13^. Moreover, high entropy and spatial coverage may denote increased variability and information richness ^42^. In this sense, the grid to place relation based on ultraslow oscillations could be beneficial for the creation of new spatial and episodic memories encoded as place cells ^15,26,27,43^.

We repeated the previous experiment, but disabling path integration during the second stage (**Suppl. Fig. S9E**). The idea behind this configuration was to isolate the effect of the ultraslow oscillation and avoid the intrinsic drift of path integration as much as possible. Hebbian learning was enabled during the first and final stages of the experiment. At the end of each learning stage, the synaptic weights connecting grid and place cells were frozen, and the activity map of each place cell was computed. When learning was turned off after the first stage, the network failed to adapt to subsequent changes in grid cell dynamics and place fields did not emerge. In contrast, during the second stage, no differences were observed between learning enabled or disabled, suggesting that during periods of oscillatory activity learning is invariant or not present ^44^.

After learning, place cells exhibited a behavior similar to grid cells: their place fields shifted over time, mirroring grid cell phase changes (**Fig. 3E**). In the control network, all place field experimented a continuous neural drift ^27,45,46^ in the same direction, caused by the intrinsic position estimation error of grid cells (**Suppl. Fig. S10**). In the oscillatory network, place fields disappeared completely during the second stage, to later reemerge at a new location once the oscillation was turned off again. Some neurons underwent complete relocation, while others retained their original field locations (**Suppl. Fig. S11**). This phenomenon closely resembled place cell remapping observed experimentally ^34,47,48^. During the oscillatory stage the activity of some place cells resembled fragments of the trajectory traversed during exploration, but only when ultraslow oscillations were present (**Fig. 3E**; **Suppl. Fig. S11**). This singularity could indicate that ultraslow oscillations promoted the transient reactivation of partial representations of recent experiences.

We quantified the spatial properties of place cell activity. The mean activity of the total population of place cells revealed that place fields were evenly distributed across the arena, with no major changes before and after the oscillation period (**Suppl. Fig. S9F**). Spatial information was strongly degraded during oscillatory periods (Friedman test, *p* < 0.001), but no significant differences were detected within the first and third stages (Dunn’s multiple test, *p* > 0.05; **Fig. 3F**). Compared with the control case, the oscillatory network differed significantly only during the second stage when oscillations were active (Wilcoxon test, *p* < 0.001, **Suppl. Fig. S9G**). A small increase in the spatial coverage was found after the oscillation period (Wilcoxon test, 0.001 < *p* < 0.01; **Fig. 3G**), but no significant differences were found between control and oscillation despite the increased variance during the second stage of the simulation (Dunn’s multiple test, *p* > 0.05, **Suppl. Fig. S9H**). Overall, aside from the observed drift, no major alterations in the spatial distribution of the place cells were determined.

We then quantified place field drift by computing the centroid of each place cell’s firing location before and after the oscillatory period. The mean drift distance was significantly larger in the oscillatory network than in the control condition (Wilcoxon test, *p* < 0.001), with no major increment in variability (Pitman-Morgan test, *p* > 0.05; **Fig. 4A**). While the drift experienced by the control network was always in the same direction (∼0°), the oscillatory network exhibited a broader distribution of drift directions (Pitman–Morgan test, *p* < 0.001; **Fig. 4B**). Population vector correlations (PVC) between pre- and post-oscillation activity maps remained above zero in both cases (one-sample Wilcoxon test, *p* < 0.001), but differed significantly between conditions (Wilcoxon test, *p* < 0.001; **Suppl. Fig. S9I**).

**Figure 4.**
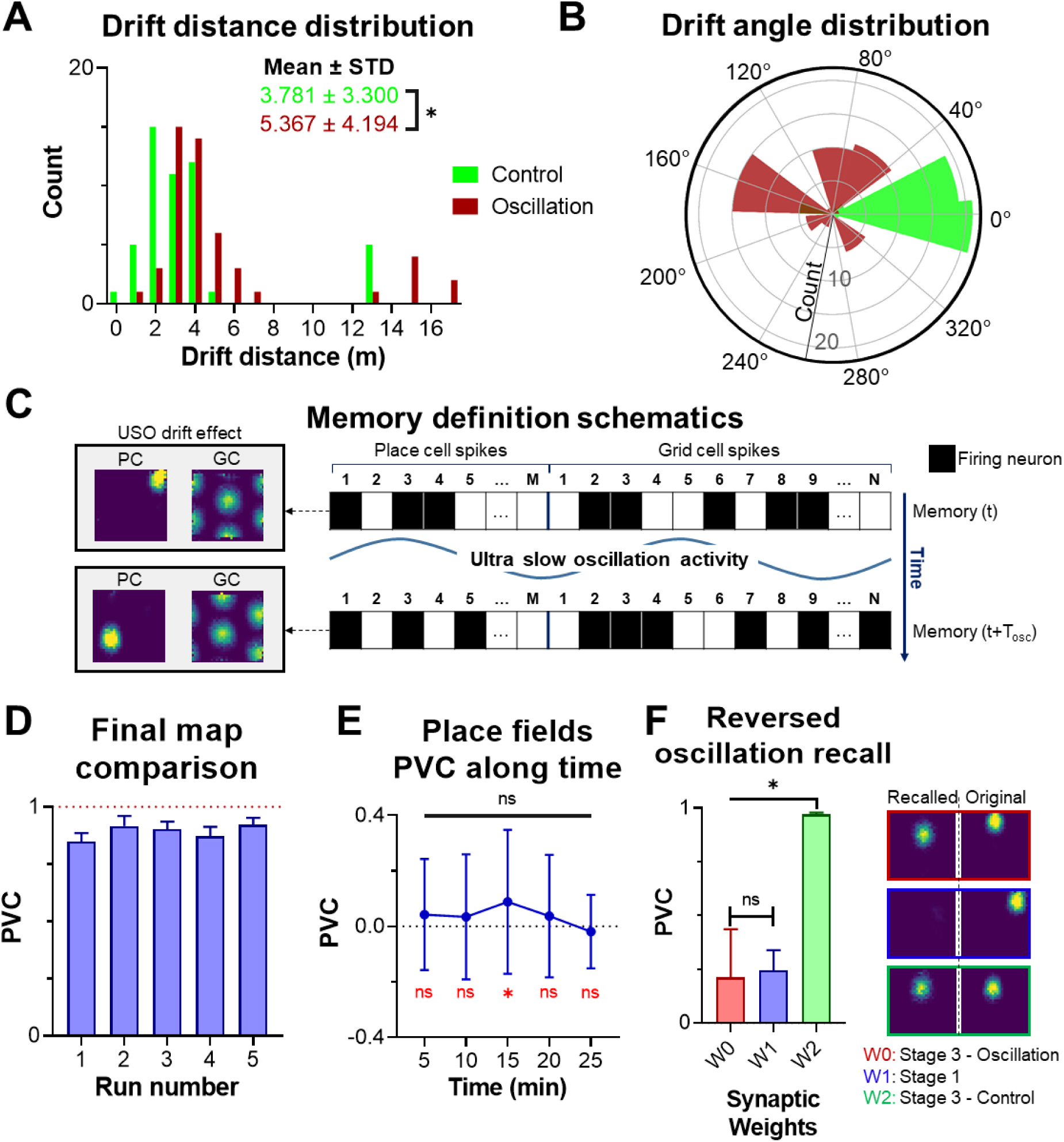
Ultraslow oscillations enable the dynamic recall of different internal representations of the same space. **A.** Drift distance comparison between control and oscillation conditions. The mean but not the variability of the oscillation condition was higher than in the control condition (Wilcoxon test, *p* < 0.001; Pitman-Morgan test, *p* > 0.05). **B.** Drift direction comparison between control and oscillation conditions. The oscillation condition exhibited a larger variability of drift directions after the remapping (Pitman-Morgan test, *p* < 0.001). **C.** Schematics of how memory ensembles were defined. In each step on the simulation, the place and grid activity was filtered, creating a vector of spikes (black squares). The combination of the grid and place vector is considered a memory ensemble. Activity maps on the left of the scheme show how the changes in the neuron firings relate to the observed drift in grid and place fields. **D.** Position recall comparison using several simulations with different trajectories but the same ultraslow oscillation characteristics. All five iteration showed a high mean PVC with low variability, indicating that the final place field distributions across all test were similar. **E.** Position recall comparison with varying length of the oscillatory stage. For each test, the place field distribution was correlated with the original 30-minutes’ simulation. Most of tje cases showed minimum PVC, with near 0 mean correlation (One-sample t-test, *p* < 0.05, red values), except for the 15-minutes experiment’s correlation which was slightly over 0 (One-sample t-test, *p* > 0.05, red values. No particular differences were found between the PVC’s among all experiments (Friedman test, *p* < 0.001, black values). **F.** Position recall after reversed ultraslow oscillation effect. Previously trained synaptic weights were trained to test whether the same place cell activity was recalled. Significant differences were found in the mean PVC across all tested synaptic weights (Friedman test, *p* < 0.001). A more detailed analysis revealed that this difference was caused by the recall of *W2* synaptic weights (Dunn’s multiple comparisons test, *p* < 0.001), while *W0* and *W1* did not show significant correlation differences (Dunn’s multiple comparisons test, *p* > 0.05)

Together, these results indicate that ultraslow oscillations primarily altered the location of place cells rather than the spatial information they convey. The drift in grid and place cells after the oscillation period was not caused by the intrinsic path integration error, but instead originated from the ultraslow oscillation. As a consequence, the same neural population of grid and place cells encoded different locations before and after the ultraslow oscillation effect.

### Ultraslow oscillation enables the dynamic access of different internal spatial representations

Previous results showed that after the oscillatory period, grid cells and place cells experienced a significant drift (**Fig. 3B, E**), different from the neural drift produced by error accumulation during path integration (**Fig. 4A, B**). This ultraslow drift caused both neuron types to fire in locations different from the ones observed before the ultraslow oscillation appearance, without major modifications in their spatial information or firing area (**Fig. 3C, D, F, G**). The firing location of biological grid and place cells is related to the encoding of spatial and episodic memories ^43^. Based on this evidence, the ultraslow oscillation drift observed in our model could indicate the codification of new memories within the same space. Because this effect was general for the entire grid and place population, the new memories suggested the emergence of a completely new internal representation of the environment.

We first assessed the difference in encodings of each location before and after the drift effect of ultraslow oscillations by quantifying memory recall. We defined a spatial memory as a grid-place ensemble (**Fig. 4C**), corresponding to a group of neurons that co-activated at the same time in a specific location of the environment ^49–51^. Memory recall was quantified using the Jaccard index, comparing ensembles overlap before and after the ultraslow oscillation at each time step. Due to the neural drift, the Jaccard index (see Methods) was not near the maximum value in the control network (One-sample t-test, *p* < 0.001; **Suppl. Fig. S12A, B**). However, the oscillatory network recall was significantly lower than the control network recall (Wilcoxon test, *p* < 0.001; **Suppl. Fig. S12A**). This effect persisted when the Jaccard index was computed by location rather than by step (Wilcoxon test, *p* < 0.001; **Suppl. Fig. S12B**). Results further confirmed that the relocation was caused by the ultraslow oscillation rather than the error accumulation of path integration (**Suppl. Fig. S12A**), and indicated a new codification of each remembered location after the oscillatory period.

We tested if this new codification was randomly generated or could be rebuilt as long as the ultraslow oscillation characteristics remained unaltered. We ran five additional simulations in which the animal followed different trajectories, while keeping the same oscillation parameters. Spatial memory recall was assessed using PVC. Because place cell activity in the model was fully determined by grid cell input and synaptic weights remained fixed, changes in grid cell dynamics were directly reflected in place cell activity and provided a sufficient measure of spatial memory retrieval. As long as the initial network state was preserved prior to oscillation onset (**Suppl. Fig. S12C**), the retrieved place cell activity after the oscillatory period closely matched the original (**Fig. 4D**). This result indicated that despite the animal traversing different trajectories, the place cells reattached to similar positions providing the ultraslow oscillation and the initial internal representation remained the same (**Suppl. Fig. S12D**). Computing the Jaccard index in each location further confirmed this phenomenon (**Suppl. Fig. S12E**). Although the obtained Jaccard values were not high, they were not significantly different from the original values in any run (Dunn’s multiple comparison test, *p* > 0.05, **Suppl. Fig. S12E**). The low resulting values could be attributed to the grid cell error, which caused slight alterations of the grid cells dynamics, leading to differences in the firing patterns compared in the Jaccard index. Nonetheless, results indicated that the new codification of each location (i.e., new spatial memories) was not random.

We examined whether the duration of the ultraslow oscillation influenced the new spatial memories. We correlated grid activity during the oscillatory period with the initial grid cell state under the assumption that if ultraslow oscillations drive the grid bump around the toroidal manifold (see Methods), the network should periodically return near its original state. Peaks in the correlation across time revealed a full cycle duration of approximately 2 ± 0.552 minutes (**Suppl. Fig. S12F**).

However, these peaks exhibited a low mean correlation (0.18 ± 0.16), significantly different from 1 (One-sample t-test, *p* < 0.001), indicating that the realignment was not perfect. This mismatch arose because the ultraslow oscillation drift was not perfectly aligned with the toroidal geometry, requiring several laps around the manifold to reach the original position, and was further altered by cumulative neural drift produced by path integration.

To test how oscillation duration affected place cell representations, we ran several iterations of the same experiment with varied oscillation periods ranging from 5 to 25 minutes in 5-minute increments. We then computed the population vector correlation between place cell activity maps from each test and the original experiment. All tests shed similar results, with a mean correlation not different from zero (Multiple one-sample t-test, *p* > 0.05, **Fig. 4E red**), except for the 15-minute test, which showed a slightly over 0 mean (One sample t-test, 0.05 > *p* > 0.01, **Fig. 4E red**). No major differences in the mean PVC among experiments were found either (Friedman test, *p* > 0.05, **Fig. 4E**). Comparisons of drift distance across oscillation durations revealed significant drift differences between oscillation period (Two-way ANOVA, *p* < 0.001; **Suppl. Fig. S12G**). Although experiments with similar (i.e., ±5 min) oscillation periods showed slightly higher mutual correlations, none approached the maximum correlation value of 1, indicating that different oscillation periods accessed distinct internal representations (**Suppl. Fig. S12H**). This outcome was consistent with the lack of perfect cycle alignment previously reported (**Suppl. Fig. S12F**). These results not only suggested that timing is crucial for memory retrieval but also that different memories can be accessed during ultraslow oscillation periods.

We examined whether the ultraslow oscillation phase influenced memory recall. Specifically, we tested whether using the complementary oscillation angle relative to the original configuration (i.e., the *θ* parameter of the ultraslow oscillation definition) would reset the location encoding to the same as before the ultraslow oscillation effect. To test this idea at the end of the experiment, we reactivated the ultraslow oscillation but using the complementary angle and assessed place cell activity using previously trained synaptic weights (**Fig. 4F**). As expected, the synaptic weights trained after the effect of ultraslow oscillation (*W0*, third stage of the oscillation condition) show no major correlations. Despite the overall low place cell activity, residual place fields still formed, which increased the mean PVC score, but were undetectable in comparison with the maximum achievable activity of place cells (**Suppl. Fig S12I**). Using the initial synaptic weights shared by both control and oscillatory conditions (*W1*), the score was not particularly high, despite the clear formation of place fields (**Fig. 4F**). In contrast, the PVC obtained with the synaptic weights trained after the third stage of the control condition (*W2*) exhibited an almost perfect recall, reflected in highly localized and strongly correlated place cell activity (**Fig. 4F**). The main difference between *W1* and *W2* was the localization of the place fields: despite the high place cell activity in both cases, the *W2* synaptic weights accounted for the path integration error suffered during the first stage of the experiment.

Overall, ultraslow oscillations enabled the creation of new spatial memories and their subsequent retrieval while accounting for intrinsic grid cell estimation drift.

## DISCUSSIONS

The results presented in this work indicated that ultraslow oscillations could emerge from a CAN model of grid cells by tuning the synaptic weights of the network. The analyses revealed that ultraslow oscillations and path integration could be seen as two independent processes that coexist in the grid cell network. During periods of path integration, ultraslow oscillations were undetectable, but their presence still severely degraded position estimation. The accumulation of error in the estimated position was caused by a drift of the grid cell dynamics originating from the ultraslow oscillation. This ultraslow drift was different from the neural drift caused by the intrinsic path integration error, and was represented as a shift of grid and place fields in space. The relocation of firing fields in both neuron types persisted despite the deactivation of ultraslow oscillation, indicating that new spatial memories emerged as a consequence of ultraslow oscillations. This ultraslow drift affected the whole grid and place population, creating a new internal representation of the environment. This new internal representation depended on the timing and phase of the ultraslow oscillation.

### Experimental vs modelled slow oscillations

The experiments in this study were inspired by two key aspects observed in the medial entorhinal cortex of mice ^16^: (1) ultraslow oscillations appeared despite the absence of sensorial stimuli, and (2) the oscillations did not depend on animal movement. We showed how altering the synaptic weights within a CAN could naturally generate ultraslow oscillations without the need for external oscillatory input or movement. In this respect, our results support the CAN model relevance ^13,29,31,35^ by proposing that ultraslow oscillation can be replicated in this type of model.

The temporal scale of the ultraslow oscillations did not match that of path integration, one of the known canonical functions attributed to grid cells ^11,17,18^. Despite ultraslow oscillations being overridden by path integration dynamics, they significantly increased position estimation error and did not contribute to error resetting. In contrast, the lasting changes induced by the ultraslow oscillation produced a neural drift, which resulted in the formation of new spatial memories. The process involved both grid and place cells, effectively creating a completely new internal representation of the same environment. These internal encodings could be recalled dynamically by controlling the phase and timing of the ultraslow drift. The high sensitivity of the slow drift to time and phase of the ultraslow oscillation may explain why ultraslow, rather than fast oscillations, are required. Ultraslow oscillations allowed the controlled modulation of the grid-place transition between spatial memories, whereas faster dynamics would induce abrupt and unreliable alterations between internal codifications. This idea goes against the MEC temporal-spatial scaffold hypothesis ^16,43^ and instead propose that ultraslow oscillation may work as a mechanism for flexibly updating the cognitive map in search of a more suitable internal representation of the environment.

### The adaptability properties of the cognitive map

Although ultraslow oscillations did not directly improve path integration in our model, their role in shaping spatial memory appears to be critical for the adaptability of the cognitive map. In these analyses, we only focus on the grid to place projection, but feedback from place cells may play a critical regulatory role. Environmental information provided by place cells could modulate the onset and termination of ultraslow oscillations, initiating them when sensory input is unavailable and suppressing them once reliable spatial cues are detected. Gonzalo-Cogno and colleagues demonstrated that ultraslow oscillations could span periods ranging from seconds to minutes, but there were not any particular stimuli that could be associated with the beginning or end of the ultraslow oscillation. We hypothesize that the lack of environmental information in their experiments could have prompted the start of the ultraslow oscillation period in search of a more suitable mental map. In darkness, the ultraslow oscillation caused the mental map to drift in a controlled manner, facilitating the learning of new grid and place associations. But these new associations depended solely on the changes in the grid cell dynamics. Meanwhile, in the presence of environmental cues, place cell activity would be prompted, which could lead to faster learning of new associations and to an early end of the ultraslow oscillation period. This behavior would result in the ultraslow oscillation not being as noticeable as in darkness, probably with shorter periods of oscillatory activity in the presence of sensorial input.

A prevalent point in our study is that ultraslow oscillations seemed to mark a turning point in our interpretation of the grid cell role in navigation. The presence of such ordered, long-timescale dynamics does not align easily with existing theoretical frameworks, creating a tension between classical models of the cognitive map and recent experimental findings. Our work alleviates some of this tension not only by showing how this oscillatory behavior can be replicated in established grid cell network models, but also by revealing their indirect but potentially crucial contribution to cognitive map formation. Rather than supporting navigation directly, ultraslow oscillations may enable the flexible reorganization and renewal of spatial representations, laying the groundwork for future experimental and theoretical investigations into their functional role.

## METHODS

### Grid cell model

The grid cell model consisted of 5 grid modules, each containing 400 neurons arranged in a toroidal neural space, represented by a *N*_*x*_ × *N*_*y*_ matrix, where *N*_*x*_ = *N*_*y*_ = 20. The position of each neuron *c* in the neural space was denoted by (*c*_*x*_, *c*_*y*_). Columns of neurons were separated by a distance *d*, and rows were separated by a distance of *d*\*√3/2. To form the characteristic hexagonal lattice of grid cells (see **Suppl. Fig. S13**), even-numbered rows were shifted horizontally by 0.5\**d*.

Within each grid module, all neurons were interconnected: each neuron excited nearby cells and inhibited distant ones. Neurons on the edge of the neural space were considered neighbors of those on the opposite edge, creating periodic boundaries. This organization, combined with local excitation, allowed neurons in the last row to excite neurons in the first row (and vice versa). Neurons from the first and last columns behaved equally. The resulting topology supported a continuous attractor dynamic on a toroidal surface, which created an activity packet that could move continuously across the neural space, following the simulated animal’s movements.

The synaptic weight connecting two cells *c*_*i*_ and *c_j_* (with *i, j* ɛ {1,2,…, *N*}) was modulated by a Gaussian function given by **Eq. 1**:

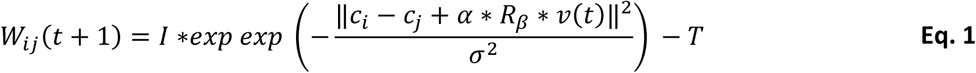

Here *I*, *T*, and *σ* were parameters controlling the amplitude, width, and offset of the Gaussian, respectively. The double bars (‖ · ‖) represented the Euclidean distance between cells *c*_*i*_ and *c*_*j*_. The planar velocity of the simulated animal, given by *v*(*t*) = (*v*_*x*_, *v*_*y*_), was the input of the network that allowed the activity packet to move around the neural space. It was modified by the gain parameter (*α*) and the rotation matrix with angle *β* (*R*_*β*_) to simulate the different scales and orientations of grid cells.

The input to any given neuron *i* at time *t* + 1 can be summarized by **Eq. 2**:

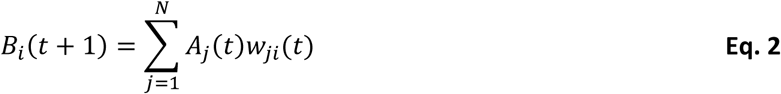

Where *B*_*i*_(*t* + 1) was a function representing the inputs to neuron *i* from all the other neurons *j* (with *j* ɛ {1, 2,…, *N*}), including itself. *A*_*j*_(*t*) was the state of neuron *j* at time *t* and *w*_*ji*_ the synaptic connection between neuron *j* and *i*, described in **Eq. 1**. The updated activity of neuron *i* at time *t* + 1 was then calculated as:

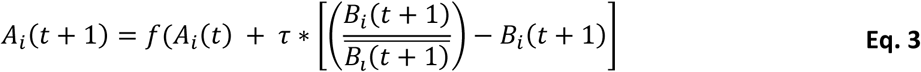

Where *τ* = 0.9 was the stabilization parameter that regulated how much the inputs affected the current state of the neuron *i* (*A*_*i*_(*t*)). The function *f* simply represented a rectification function that limited the activity of any cell between 0 and 1.

The network was initialized with random values between 0 and 1/√*N*. Before any simulation, the network was first executed with no inputs for 100 iterations, until it reached a stable point. Once this initialization was done, a clear activity packet emerged in each of the grid modules. The location of the initial bump was random and varied across simulations. A velocity signal was used as input to move the activity packet around the neural space. The velocity vector *v*(*t*) at each time-step was random, with a mean velocity of 0.1 ± 0.05 m/s.

### Grid scale and gain relation

The simulations used five grid cell modules, each with a different gain parameter linearly spaced between 0.02 and 0.12. All modules shared the same orientation. This configuration was chosen to avoid distortions in the decoded trajectories during path integration. In the model, simulating different grid orientations required rotating the input velocity vector. This mechanism implied that during path integration, the decoded vector had to be “unrotated”, which could lead to deformations in the predicted trajectory.

The gain parameter was not the same as the scale of the grid cell. Using non-linear least square we determined that the relation between grid and scale was given by **Eq. 4**

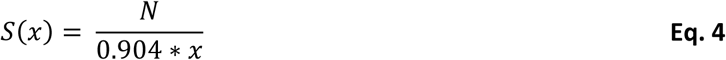

Where *S*(*x*) was the scale of the grid module with gain *x* and *N* the number of grid neurons around the *x*-axis of the torus. The fit yielded a mean squared residual of 0.00019 and a maximum covariance of 4.61 * 10^-8^. This function was used to estimate the spatial scale of each module from its gain, which was then utilized for decoding position during path integration.

### Ultraslow oscillation

Ultraslow oscillations were generated by defining a directional path within each grid cell module that induced a slow drift of the activity packet around the toroidal neural space. The path was parameterized by two values *ρ* and *θ*. The parameter *ρ* controlled the strength of the drift. The larger *ρ* value was, the stronger the directional bias and the faster the activity packet moved along the path. The angle *θ* ɛ [-*π*, *π*] indicated the orientation of the path. For example, an angle of *θ* = 0 implied that the path would travel around the torus parallel to the *x*-axis, while an angle of *θ* = *π*/2 meant the path wrapped around the torus parallel to the *y*-axis.

This directional path was implemented by introducing small, asymmetric perturbations to the synaptic weights of the grid cell network. Normally, the synaptic weights on the network were stable and symmetrical in order to create a single attractor. By inducing a small guided perturbation on those synaptic weights, a directional “pull” was introduced, causing the attractor to drift. From a given neuron *c*_*i*_ in the network, located in (*c*_*ix*_, *c*_*iy*_), the next two target neurons along the path were *c*_*j*_ and *c*_*k*_, positioned in (*c*_*ix*_+*d*_*x*_, *c*_*iy*_) and (*c*_*ix*_, *c*_*iy*_+*d*_*y*_) respectively. The direction of flow, represented by *d* = (*d*_*x*_, *d*_*y*_), was computed based on the angle *θ* as follows:

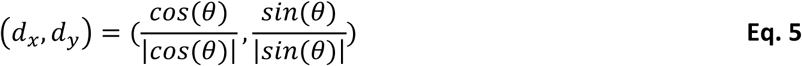

Where *d*_*x*_, *d*_*y*_ ɛ {-1, 0, 1} indicated the discrete direction of flow along the *x* and *y* axes. The weight connecting cells *c*_*i*_ and *c*_*j*_ (*w*_*ij*_) was then modified by |*cos*(*θ*)| * *ρ*, and the weight connecting cells *c*_*i*_ and *c*_*k*_ (*w*_*ik*_) was modified by |*sin*(*θ*)| * *ρ*.

This approach took into account that neurons were arranged in a matrix, but their true physical layout was hexagonal, with some cells offset horizontally to achieve the triangular tessellation. Because of this organization, diagonal neighbors in the matrix might not be directly adjacent in physical space. Therefore, the perturbation was applied only to neurons that were aligned along the *x* or *y* axes, ensuring consistency in the directional flow regardless of tessellation offset. This process was repeated for all neurons in each module. The ultraslow oscillation parameters *ρ* and *θ* could differ across modules or remain the same, depending on the simulation setup.

### Place cell generation

Place cell generation was implemented using a simple Hebbian learning rule with competition among grid cells and place cells. For both neuron types, the winners were selected using k-Winner-Take-All (k-WTA), as it provides a computationally efficient approximation of the cortical inhibitory competition mechanism ^52,53^. For place cells, a single winner was selected from the population, whereas for grid cells, two winners were chosen per module.

Competition among grid cells was implemented at the module level, as place cell formation benefits from the variability in scale of grid cells ^54^. Because grid cells within a module shared the same scale and rotation, selecting all competition winners from a single module would cause the resulting place cell to inherit the firing pattern of that module. By forcing grid cells from different modules to participate in the synaptic connections, each place cell developed a single, stable place field. This modular competition mechanism has been extensively discussed in the literature ^31,55,56^.

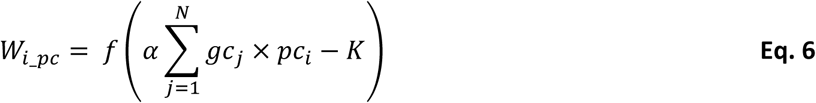

Here *W*_*i*_*pc*_ denoted the incoming synaptic connections of place cell *i* ɛ {1,2,…*M*} with *M* being the size of the place cell population. The term *gc*_*j*_ represented the activity of grid cell *j* ɛ {1,2,…*N*} with *N* being the total number of grid cells, while *pc*_*i*_ was the activity of place cell *i*. The learning rate *α* = 10^-2^ and the decay constant *K* = 10^-3^ regulated synaptic growth by avoiding over-fitting and suppressing connections between asynchronously firing neurons. The *f* function referred to a simple rectification and normalization function that constrained synaptic weights to the range [0,1]. Normalization was applied modularly to prevent a single module from dominating the synaptic connectivity.

All experiments were performed in the dark with no sensorial stimuli present, meaning that place cell activity was solely determined by grid cell activity. Because weights were rectified, inhibitory connections were eliminated, and place cell activity became noisy. To mitigate this effect and create clearer place fields, place cell activity at each time step of the simulations was computed as:

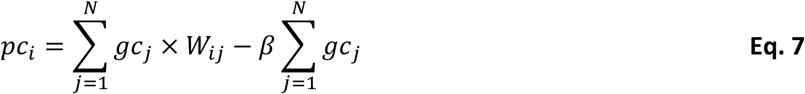

Where *pc*_*i*_ and *gc*_*j*_ represented the activity of place cell *i* and grid cell j, respectively, while *W*_*ij*_ was the synaptic weights connecting both of them. The second term of the equation introduced a global inhibitory drive proportional to total grid cell activity, regulated by a constant *β* = 10^-3^ × *G*, with *G* being the number of grid modules. This term captured the effect of hippocampal interneurons ^57^ and promoted the emergence of single, well-defined place fields.

### Data analysis

#### Cell filtering and selection of oscillation parameters

Gonzalo-Cogno and colleagues reported that not all medial entorhinal cortex grid cells participated in the observed oscillations. To account for this phenomenon in our model, we filtered the simulated neuron population to retain only those cells that exhibited oscillatory activity. Following the procedure in ^16^, we defined for each neuron an activation threshold equal to its mean activity over time plus 1.5 times its standard deviation. In our model, all cells shared the same activity distribution, differing only by a spatial offset. Therefore, instead of using individual thresholds, we used a single global threshold equal to the maximum of all neuron-specific thresholds. This threshold was applied to detect spikes, and any neuron that did not spike at least once during the simulation was discarded. The remaining cells were used for all subsequent analyses.

To find the path parameters that induced similar oscillatory frequencies, we systematically explored the oscillation parameter space. First, we ran multiple 30-minute simulations with no animal movement, sampled at 7.73 Hz (13,914 samples total). The purpose behind this procedure was to avoid any kind of interference produced by path integration. The resulting activity of each simulation was filtered and normalized to the range [0, 1] using the maximum activity of each cell. Because all neurons behave equally, measuring the oscillatory frequency of a single cell was sufficient to determine the frequency of the entire module. For each module, we selected one representative neuron and computed its autocorrelogram. We then estimated the power spectral density (PSD) using Welch’s method from the Scipy library, with a Hamming window equal to the recording length and 50% overlap between consecutive windows. Peaks in the PSD with amplitude greater than 1 were detected using Scipy’s *find_peaks* function. This analysis was repeated for all combinations of oscillation parameters *θ* and *ρ*, with *θ* ɛ [-*π*, *π*] and *ρ* ɛ [0.01, 5.0]. The combination pairs were then grouped according to the frequency of their PSD peaks. From each frequency group, a subset of parameter pairs with size equal to the number of grid modules was selected for the final simulations.

### Stacked Z-score

The stacked z-score correlation was calculated to assess whether all neurons from the network are firing at the same frequency. After filtering and normalizing the neural activity, the z-score was calculated from the autocorrelation of each cell individually. The autocorrelation was used to calculate the PSD, using the Welch method, with a Hamming window of 8192 samples (or 17.6 minutes at a sampling rate of 7.73 Hz) and 50% overlap. The z-scores were then normalized and stacked into a matrix, sorting them based on the maximum value reached by their PSD. A Gaussian filter with a kernel standard deviation of 10 was applied to the resulting matrix to reduce the noise produced by harmonic frequencies too similar to each other.

### Autocorrelation cell sorting

To visualize ultraslow oscillations in raster plots, we established an ordering criterion based on cross-correlating neural activity over time. This approach identified near-instantaneous co-firing between neurons.

First, the neural population was filtered and its activity normalized to the range [0, 1] using the maximum activity of each cell (see Methods: *Cell filtering and selection of oscillation parameters*). We then constructed an *N* × *N* correlation matrix *R*, where *N* was the number of filtered neurons that could potentially contribute to the oscillation. For each neuron pair (*i, j*), their activity traces were cross-correlated using a maximum time lag of 39 bins (5 seconds at a sampling rate of 7.73 Hz). This time window ensures that cells firing within a similar timeframe were strongly correlated.

From each cross-correlation, we extracted the maximum value and multiplied it by the sign of the corresponding time lag peak. This last step was done to keep the firing order, as a negative correlation implied that neuron *i* fired before neuron *j* and vice versa. The result from this operation was then stored in the matrix as *R*_*i*j_. At the same time, the reciprocal value (*R_ji_*) was also calculated as *R*_*ij*_ and stored in its respective index in the matrix.

Once the *R* matrix was completed, the maximum value was determined, and its associated *i*-cell was taken as the seed cell. The rest of the neurons were organized in a descending manner based on their correlation with the seed cell. Finally, the firing peaks of each neuron were stored in a matrix of *N* × *T* with *T* being the number of samples of the simulation. The rows of the matrix were ordered based on the previously calculated criterion.

### PCA cell sorting

The second order criterion was based on principal component analysis (PCA). After filtering and normalizing the neural population activity, the PCA method (Scipy’s implementation) was applied to the activity matrix, with neurons as rows and time steps as columns. From this procedure, we extracted the first two principal components (*PC*_1_ and *PC*_2_), defining a new two-dimensional space (*S*_*PCA*_) in which the activity of the neurons could be projected. Only two components were retained, as this was the minimum required to capture a one-dimensional dynamic such as the observed ultraslow oscillation.

Each neuron was represented in *S*_*PCA*_ by its PCA scores, forming a two-dimensional vector. The angle of this vector with respect to the *PC*_1_ axis was calculated as:

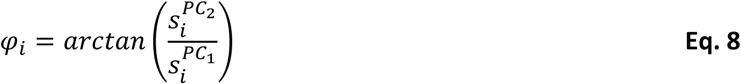

Where *φ*_*i*_ ɛ [-*π*, *π*] was the angle formed by neuron *i* projected in *S*_*PCA*_, and **s*_i_^*PC*j^* indicated the score of neuron *i* along each of the *PC*_*j*_. The angle *φ* was then used to order the firing peaks of all neurons in a *N* × *T* matrix, analogous to the cross-correlation method. Unless otherwise specified, the PCA-based criterion was used throughout this study due to its computational efficiency.

### Oscillation period and frequency

To estimate the oscillation period and frequency, we applied PCA to the neural activity with time steps as rows and neurons as columns. From this analysis, the first two principal components (*PC*′_1_ and *PC*′_2_) were extracted, defining a new two-dimensional space (*S*_*osc*_). The state of the neural population at each time (*A*(*t*)) was projected in this space, resulting in the vector (*S*_*osc*1_(*t*), *S*_*osc*2_(*t*)), obtained by taking the dot product of *A*(*t*) with the eigenvectors of *PC*′_1_ and *PC*′_2_, respectively. This representation reduced the dimensionality of the network while preserving its population dynamics.

The phase of the oscillation at each time step was calculated as:

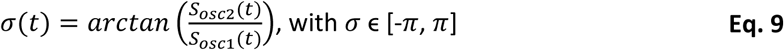

The oscillation amplitude at each time step was defined as *sin*(*σ*(*t*)). A PSD analysis was then applied to this signal, using the Welch’s method with a Hamming window of 8192 samples (17.6 minutes at a 7.73 Hz sample frequency) and 50% overlap. The dominant frequency *f*_*max*_ was identified as the maximum peak of the PSD.

For *f*_*max*_ to be considered the oscillation frequency, two conditions had to be satisfied: (1) *PSD*(*f*_*max*_) was greater than nine times the mean power of the tail of the spectrum, with the tail defined as {*f* | *f* > *f*_*max*_}) and (2) *PSD*(*f*_*max*_) was greater than nine times the minimum power of the head of the spectrum, with the head defined as {*f* | < *f*_*max*_}). If both conditions were met, the oscillation frequency was defined as *f*_*max*_ and the oscillation period as 1/*f*_*max*_. Otherwise, no oscillation was considered present, and both frequency and period were set to zero.

### Spectrograms and scalograms

Spectrograms were used to identify patterns of neural activity and detect oscillatory dynamics at certain frequencies. In this study, we utilized the Scipy implementation of the spectrogram method with a sampling frequency of 7.73 Hz, using a Tukey window with segments of 512 points (∼264 seconds) and a 50% overlap between windows. A limitation of spectrograms was found for a given parameter configuration because they were sensitive only to a restricted frequency range. This configuration allowed reliable detection of very low frequencies (< 0.01 Hz), while higher-frequency components were not captured and therefore exhibited negligible power in the spectrogram. To overcome this limitation, we additionally used scalograms, which provided a multi-scale time– frequency representation by effectively combining information across frequency bands within a single plot.

Scalograms in this paper were computed using the *cwt* function of PyWavelet. We utilized a sampling frequency of 7.73 Hz and a Morlet wavelet with a bandwidth of 1.5 and a center frequency of 1.0. The transform was computed over 300 logarithmically spaced scales, corresponding to frequencies between 0.001 Hz and 0.3 Hz, adjusted for the sampling frequency. This approach enabled simultaneous visualization of both ultraslow and slightly higher frequency oscillatory activity.

### Oscillation score

The oscillation score quantified the degree to which grid cell activity exhibited oscillatory dynamics. It integrated two ordering approaches by constructing a joint distribution of the time lags that maximized the cross-correlation between cell pairs (*τ*) and their corresponding angular distances in the PCA plane (*d*).

We first computed the phase of each neuron (*φ*_*i*_) in the PCA plane (see **Eq. 8**). This phase also served as a parameter to define alternative ordering criteria. Three criteria were evaluated: (1) using data from the first third of the experiment, during which the velocity input was disconnected; (2) using the second third, when position was estimated using the velocity signal; and (3) using the entire activity sequence. These distinct orderings allowed us to assess whether oscillatory patterns emerged preferentially during specific experimental stages.

The angular distance between each pair of cells was defined as the difference between their phases, normalized to the interval [−*π*, *π*):

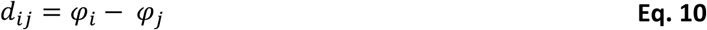

For each cell pair, we computed the cross-correlation function and extracted the time lag (*τ*) corresponding to its maximum value, with a maximum lag of 248 s or 1917 samples at 7.73 Hz.

The resulting angular distances and time lags were assembled into a two-dimensional histogram. The phase range [−*π*, *π*) was divided into 11 bins (∼0.57 rad each), and the time lag range [0, 248) s into 96 bins (∼5 s each). The histogram was normalized by the total number of cell pairs, yielding the joint distribution of *d* and *τ*.

For each row of the 2D histogram (i.e., for each *d* bin), we computed the power spectral density (PSD) using the Welch method with a Hamming window of 128 bins and 50% overlap. A PSD peak was considered significant if its amplitude exceeded both (1) ten times the mean power in the tail (from the peak frequency to the highest frequency) and (2) 4.5 times the mean power in the head (from 0 Hz to the peak frequency). When both conditions were met, the corresponding *d*_*i*_ bin was classified as oscillatory.

The oscillation score was then defined as the proportion of *d* bins in which oscillatory behavior was detected.

### Position estimation using grid cells

In our model, grid cell activity formed a localized “bump” that continuously shifted across a toroidal neural space based on a velocity input, replicating a simulated animal movement through the environment. To estimate the animal’s position, we computed the bump displacement at each time step for every grid module. This displacement represented the inferred movement in the neural space, which may not directly correspond to the true physical displacement. To map it onto real-world coordinates, the bump displacement was multiplied by the scale of each module (see Methods: *Grid scale and gain relation*) and added to the previously estimated position. By averaging the contributions of all grid modules, we obtained a single position estimate defined by Cartesian coordinates (*x*,*y*) relative to the starting point. The starting position was set at the center of the arena, corresponding to (*x*,*y*) = (0, 0).

### Drift directionality

The direction of the drift was computed using the grid cell activity from the first third of the experiment, during which the network did not receive velocity input. In this period, any displacement of the activity bump was caused solely by the ultraslow oscillation and thus followed its direction. Using the position estimation procedure described in “Position estimation using grid cells”, we reconstructed the trajectory traced by this drift. The coordinates of the last step of the trajectory were then used with the *arctan2* function from the NumPy library to calculate the drift direction.

### Activity map generation

The activity map for both grid cells and place cells was generated using the *binned_2d_statistic* method from the Scipy library. It divided the arena into *B*=30 bins of ∼0.06 m^2^. For each step of the simulation, the neural activity was assigned to one of the bins based on the (*x*,*y*) position reached at that time of the simulation. Then the activity in all bins was averaged and saved as the final value of the bin. This process was repeated for each neuron. The result was a matrix of *B* × *B* bins showing the spatial behavior of the cell.

#### Spatial analysis

Spatial metrics were used to quantify the relationship between neural activity and physical space. All measures were derived from the activity map of the cell, and capture complementary properties of the cell, such as its phase, the amount of spatial information it conveys, or the percentage of the arena that it covers while active.

#### Phase distance

This measure quantified the displacement of a grid cell’s firing fields relative to their original location. It was calculated using the two-dimensional cross-correlation between the activity map at a given stage and the reference activity map from the initial stage. Simulations were divided into three stages, with the trajectory associated with each stage being long enough to cover all the enclosure. The activity map was computed for each of those stages and cross-correlated using the *correlate2d* method from the Scipy library. The peak location of the resulting cross-correlogram indicated the spatial offset where both activity maps match. Identical representations yield a peak at (0,0), whereas displaced representations produce a shifted peak. By calculating the Euclidean distance of the peak to the center using its matrix subindexes, we obtained the phase drift between the initial and final stages of the simulation.

#### Spatial information

It measured the specificity of the firing fields of a neuron, based on how their activity was distributed around space (Markus et al., 1994; Skaggs et al., 1992). It was calculated using the activity map of the cell, dividing the arena into *B*=30 bins of ∼0.06 m^2^ (see Methods: *Activity map generation*). The spatial information of the cell was computed as:

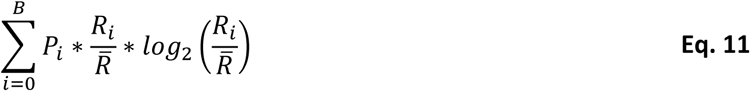

Where *i* ɛ {1,2,…,*B*} with *B* being the total number of bins. *P*_*i*_ was the probability of occupying bin *i*, calculated as the total number of steps of the trajectory between the limits of the bin divided by the total number of steps. *R*_*i*_ was the mean cell activity in bin *i*, while *R̅* is the overall mean activity.

#### Spatial entropy

This measure provided a complementary characterization of spatial information, indicating how spread out the firing fields of a cell were. It was computed using Shannon’s entropy formulation (Shannon, 1948):

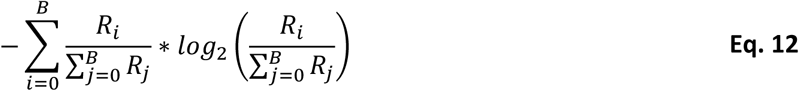

Where *R*_*i*_ and *R*_*j*_ were the mean cell activity in bin *i* and j, respectively, with *i*, *j* ɛ {1,2,…,*B*} and *B* being the total number of bins.

#### The effective spatial coverage

(ESC). The fraction of the arena in which a neuron was considered active. A null distribution was created using the mean activity from the shuffled bins of the neurons’ activity maps. A minimum activity threshold to consider the cell active in a bin was selected from a set of candidates using a Benjamini-Hochberg False Discovery Rate (**Eq. 13**)

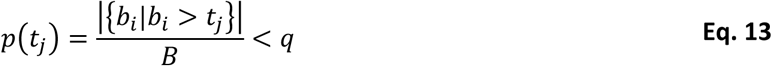

Where *p*(*t*_*j*_) was the *p*-value associated with the threshold candidate 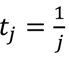 with *j* ɛ [1;10]. The threshold values range from (0;1] based on the maximum and minimum neural activity. The mean neural activity of bin *i* was represented as *b*_*i*_, with *i* ɛ {1,2,…,*B*} and *B* being the total number of bins. The symbol |.| indicated cardinality of the set. The *q* value indicated the maximum expected rate of false discoveries, which was set to 0.05. This procedure ensures that no more than 5% of bins classified as active were expected to be false positives.

Once the final threshold was obtained, the effective spatial coverage of neuron *c* was computed as:

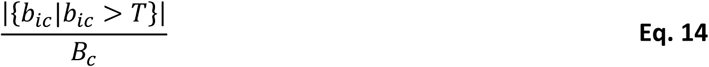

Where *b*_*ic*_ was the mean activity of bin *i* from cell’s *c* rate map, and *T* was the final neural activity threshold calculated. The symbol |.| indicates cardinality of the set.

### Neural ensembles analysis

Neural ensemble metrics were used to assess the memory recall capacity of the model. Memories can be understood as a fixed set of neurons or neural ensemble that consistently coactivate in response to the same stimulus. Because no sensory input (e.g., vision) was available in our experiments, neural firing depended solely on the position of the simulated animal, conveyed through grid cells. Accordingly, the simultaneous activation of grid cells and place cells at a particular moment of the simulation constituted a memory ensemble. Memory ensembles were extracted by using a threshold over the neuronal activity according to *γ*: {*x*| *A*_*x*_(*t*) > *γ*}, with *x* being the cell number and *A*_*x*_(*t*) the activity of cell *x* at time *t*. For grid cells, the threshold was set to the mean activity of each cell plus 1.5 times its standard deviation, whereas for place cells, a threshold of zero was used.

**The Jaccard similarity index** quantified the degree of similarity between two memory ensembles in a range between 0 (no overlap) and 1 (identical ensembles):

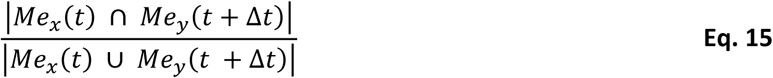

Where *Me*_*x*_(*t*) and *Me*_*y*_(*t*) referred to the memory ensembles of simulations *x* and *y* at time *t* of the simulation. If the memory ensembles were extracted from the same simulation (i.e. *x* = *y*) at the same time (i.e. Δ*t* = 0), then the Jacard index is equal to 1.

The Jacard index was computed either across time or across space. In the temporal analysis, ensembles were extracted at each time step and compared directly. In the spatial analysis, the arena was divided into *B*=30 bins, each assigned with a memory ensemble computed as the filtered activity of grid and place cells at the first moment the simulated animal entered that bin. Similarity was then assessed by comparing ensembles associated with the same spatial bin across different simulations or time points.

**The population vector correlation (PVC)** quantified the degree of similarity between two place cell populations based on their activity maps (**Eq. 16**). Because in our model the grid cells determined place cells, the PVC could be used to measure the degree of similarity between recalled memories as long as the synaptic weights connecting grid and place remained unaltered.

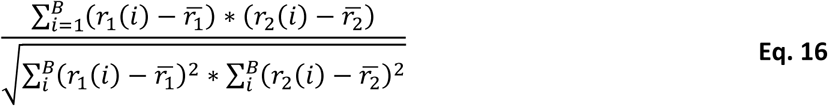

Where *r*_1_(*i*) and *r*_2_(*i*) were the ratemap value in bin *i*, with *i* ɛ {1,2,…,*B*} and *B* being the total number of bins.

## STATEMENTS AND DECLARATIONS

## Acknowledgments

Thanks to Elias Todorovich for comments on the final manuscript. To the Gonzalo-Cogno Lab for discussions on the topics reported in the latest version of the manuscript. A preliminary version of this work was presented as a poster at the SAN2025 Annual Meeting (3-5/10/2025). LS and JAF-L are supported by The National Scientific and Technical Research Council (CONICET), Argentina, through a Type-1 Doctoral Scholarship, and the Scientific and Technological Researcher Career program (DI-2019–2516-APNGRH-CONICET), respectively. MP is supported by The Comisión de Investigaciones Científicas de la Provincia de Buenos Aires (CICPBA), Argentina.

## Author Contribution

LS: Conceptualization, Investigation, Coding and Writing & Editing – original and final draft. JAF-L: Conceptualization, Funding acquisition. Supervision. Writing & Editing – original and final draft. MP: Coding of spectral analyses. Review – final draft.

## Conflicts of interest/Competing interests

The authors declare that the research was conducted without any commercial or financial relationships that could be construed as a potential conflict of interest.

## Funding

LS is supported by The National Scientific and Technical Research Council (CONICET) Argentina, through a doctoral scholarship.

JAF-L is supported by CONICET through the Scientific and Technological Researcher Career program (DI-2019–2516-APNGRH-CONICET).

## Ethics approval

This article does not contain any studies with human participants performed by the authors.

## Clinical trial registration

The authors affirm that there is not clinical trial information associated to this manuscript.

## Permission to reproduce material from other sources

Permission should be obtained in writing from the authors if all or part of the text is used in a publication.

## Code availability

The code will be publicly available at https://github.com/LabNeuroAI/USO_GC_drifting_model after publication.

## Notes

### Competing Interest Statement

The authors have declared no competing interest.

