## Supplemental Information for "Ultraslow entorhinal oscillations shape spatial memory through grid cell drifting"

Luca Sarramone (1,2,*), Matias Presso (1,4), Jose A. Fernandez-Leon (1,2,3,*)

(1) NeuroAI Lab, Fac. Cs. Exactas-INTIA, Universidad Nacional del Centro de la Provincia de Buenos Aires (UNCPBA), Tandil, Buenos Aires, Argentina
(2) Consejo Nacional de Investigaciones Científicas y Técnicas (CONICET), Buenos Aires, Argentina
(3) CIFICEN (CONICET–CICPBA-UNCPBA), CCT-Tandil, Buenos Aires, Argentina
(4) Comisión de Investigaciones Científicas de la Provincia de Buenos Aires (CIC), Buenos Aires, Argentina

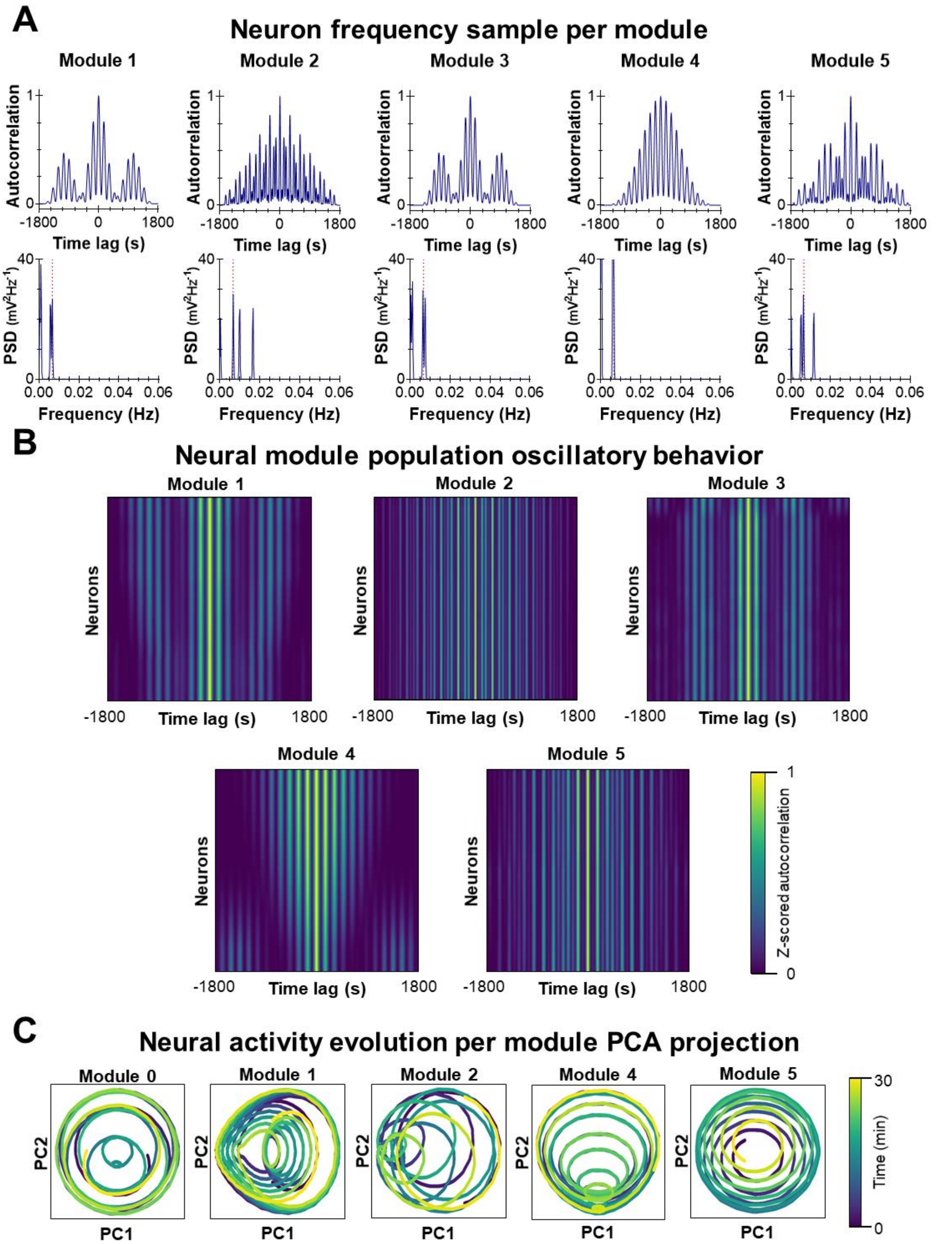


**Supplemental Figure S1.** Autocorrelation (top) of cell activity and the PSD (bottom) calculated from it. The five columns correspond to five different representative neurons from each module. The dotted line signals the frequency under which all the neurons were grouped (see Methods). **B.** Stacked z-score of individual neuron's activity correlations grouped by grid module and organized by maximum PSD activity. The pattern shows that all neurons from the same module fired at a similar frequency because they shared the same oscillation parameter values. **C.** PCA projections of the grid cell activity from each module into a 2D space. Activity exhibited an oscillatory dynamic similar to the one observed in vivo. Color indicates simulation time.

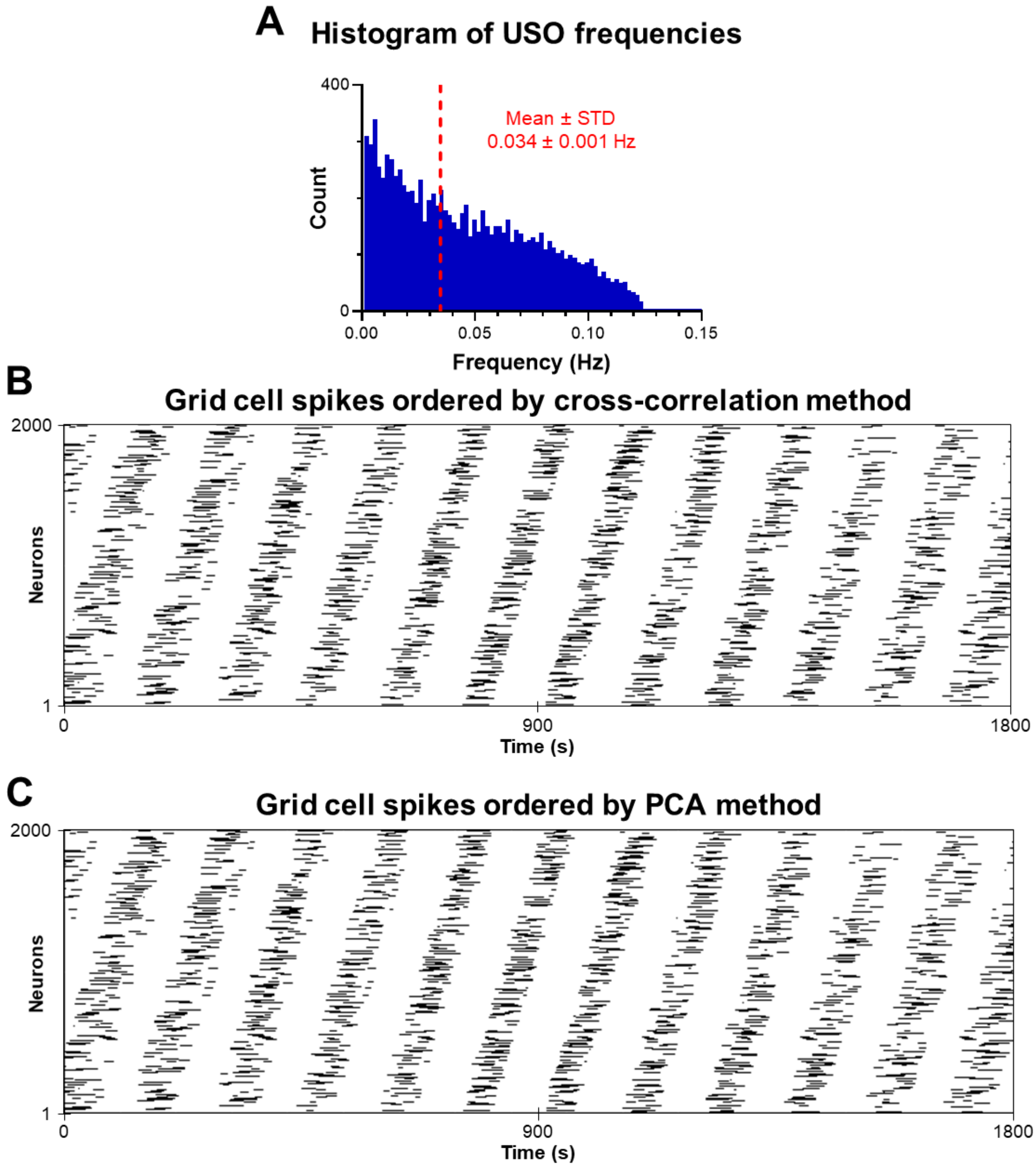


**Supplemental Figure S2. A.** Histogram of oscillation frequencies achieved by combining different oscillation parameters ($\theta$ and $\rho$) values. Read line indicates weighted mean oscillatory frequency**.** Parameter combinations with harmonics were included in the frequency bin with the largest PSD value, as well as in all the harmonics' frequency bins, to cover the maximum range of possible oscillation frequencies. **B.** Raster plot example of neural spikes over time, sorted by the cross-correlation criterion (see Methods). The simulation lasted 30 minutes (1800 s), and a total of 2000 neurons were registered. The frequency of this oscillation was 0.0066 Hz, the same as that used in the rest of the experiments in the paper. **C.** Same as B, but using the PCA method for ordering.

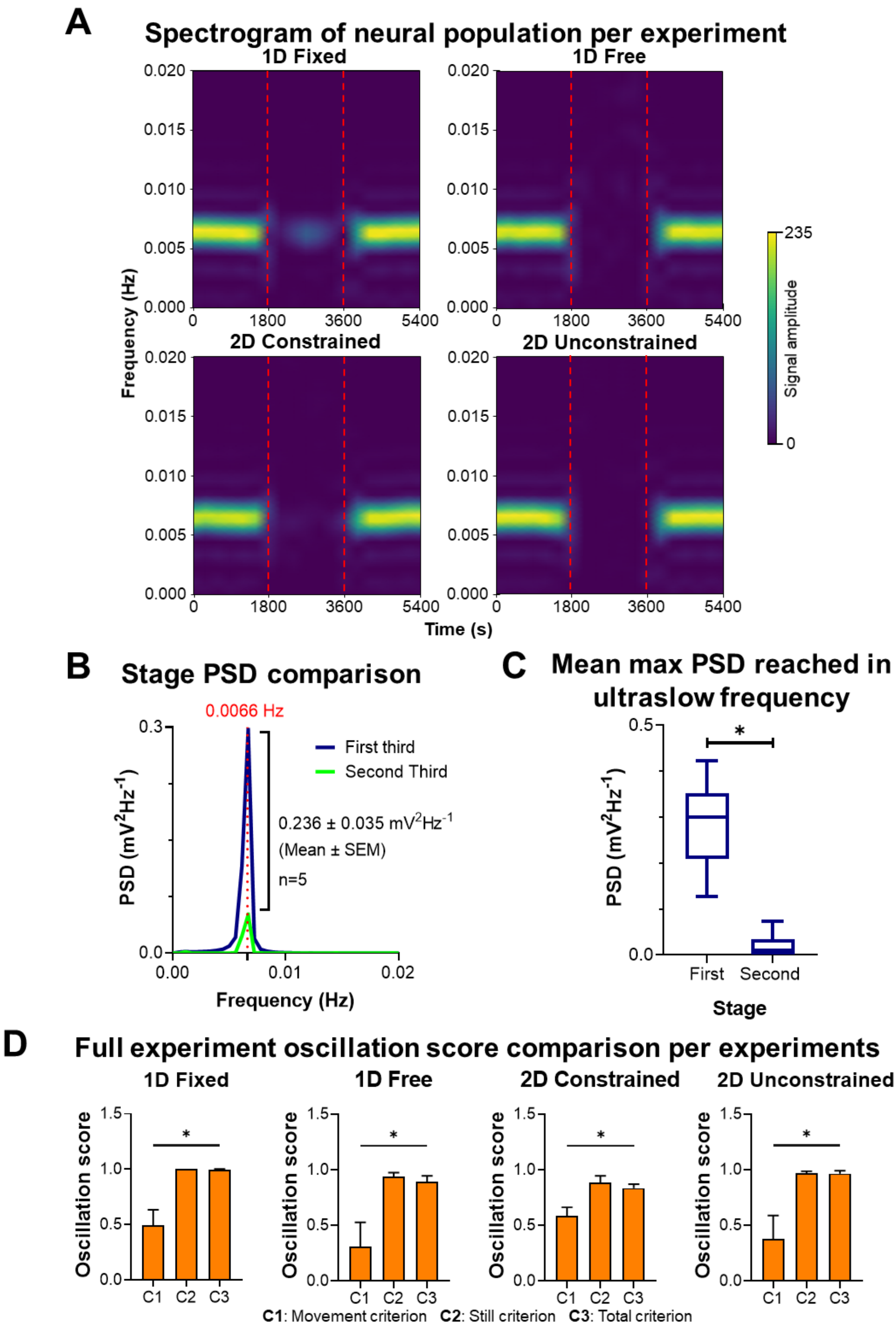


**Supplemental Figure S3. A.** Spectrogram of grid cell population activity for the four experimental conditions. Red line separates the stages of the experiment. During the first and third stage where only ultraslow oscillations were active, the neural activity concentrates around the 0.0066 Hz frequency. **B.** PSD comparison example between the first and second stage of the 1D-Fixed experimental condition (see **Supplemental Fig. S4** for other experimental conditions). The comparison is focused on the ultra-low oscillation frequency (0.0066 Hz). For each experimental condition, $n = 5$ repetitions were done, and the mean difference between the PSD peaks of each stage of the same experiment was calculated. **C.** Mean PSD peaks' power of all experiments from every condition ($n = 20$) comparison between the first and second stage. A significant difference was found (Wilcoxon test, $p < 0.001$). **D.** Oscillation scores comparison between order criteria for each experiment. Scores were calculated using the activity from the whole experiment, without regard for stage separation. Statistical differences were found among all three experiments (Friedman's test, $p < 0.05$).

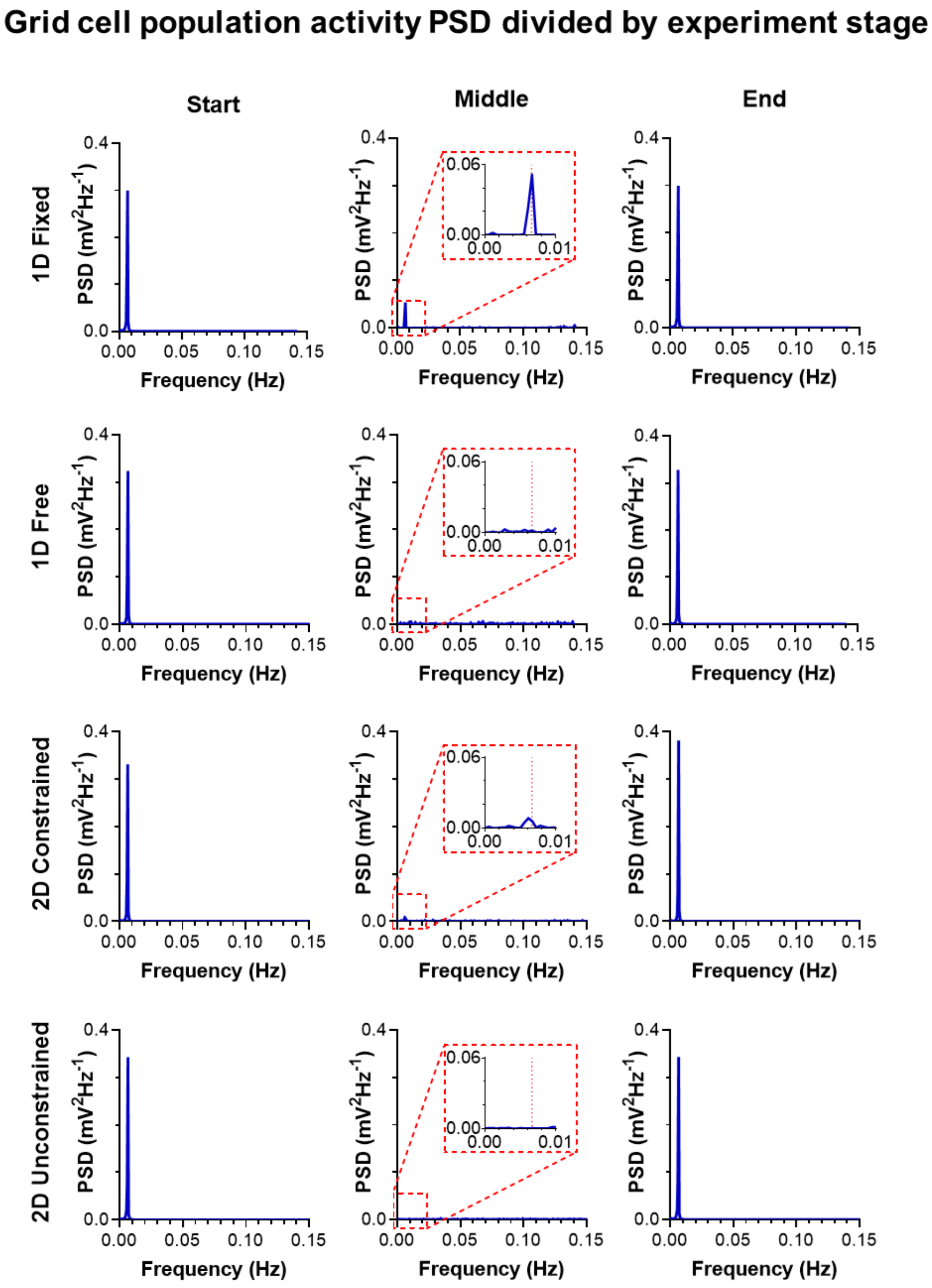


**Supplemental Figure S4.** PSD of grid cell activity separated by stage and by experimental condition. Columns indicate the stage (from left to right): first stage (start) with only ultraslow oscillation, second stage (middle) with ultraslow oscillation and path integration, and third stage (end) with only ultraslow oscillation again. Rows separate experimental conditions. The scale during the second stage was modified to adjust for the new frequency peaks, otherwise the PSD would look like a flat line.

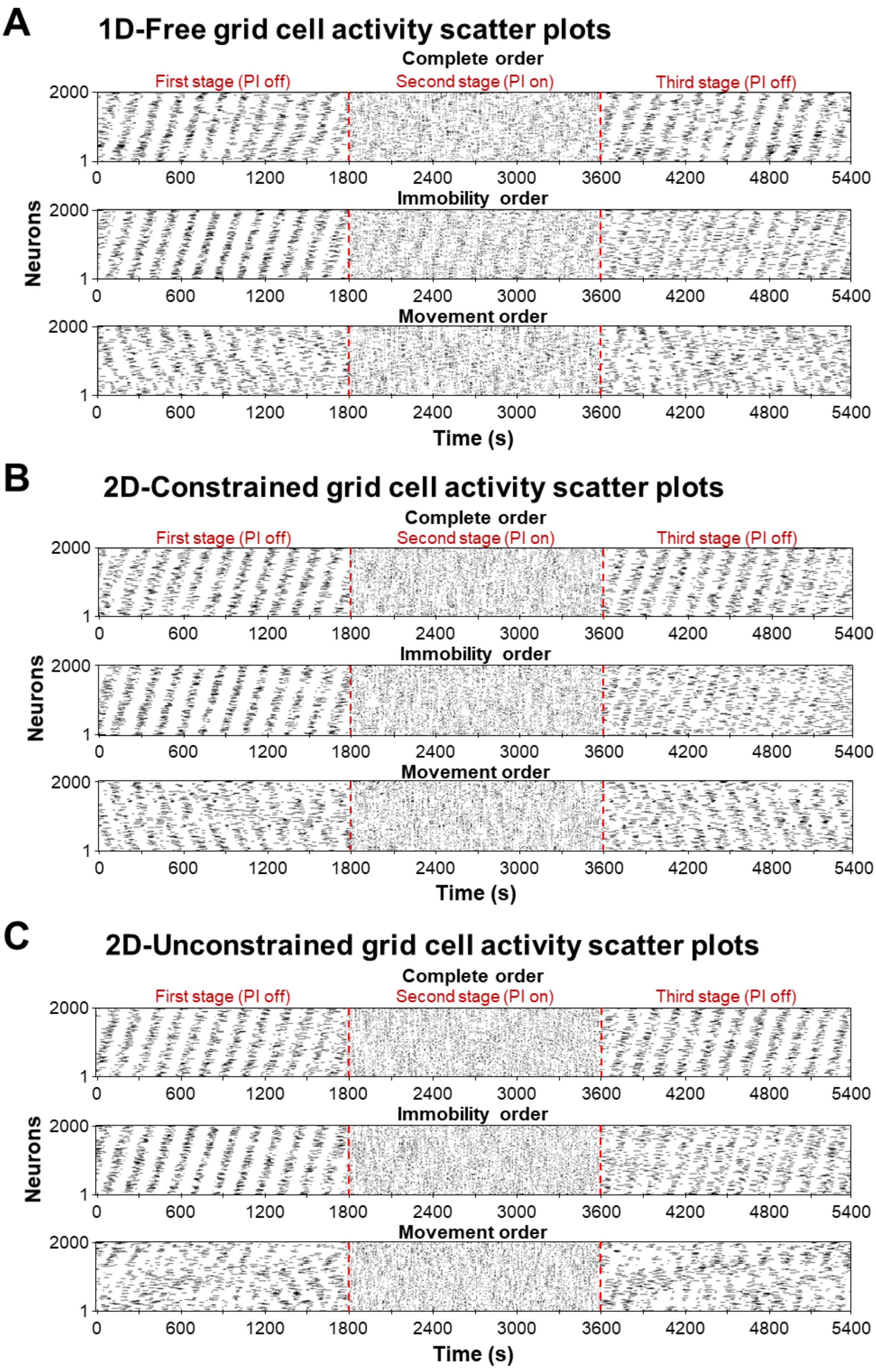


**Supplemental Figure S5.** Raster plots of neuron activity over time for the three missing experimental conditions: 1D-Fixed, 2D-Constrianed, and 2D-Unconstrained exploration. All of them were sorted by the same three different criteria explained in **Fig. 2B**. From top to bottom: (I) Based on the neuron activity through the whole experiment; (II) Based on the neuron activity of only the first third of the experiment; (III) Based on the neuron activity of only the second third of the experiment.

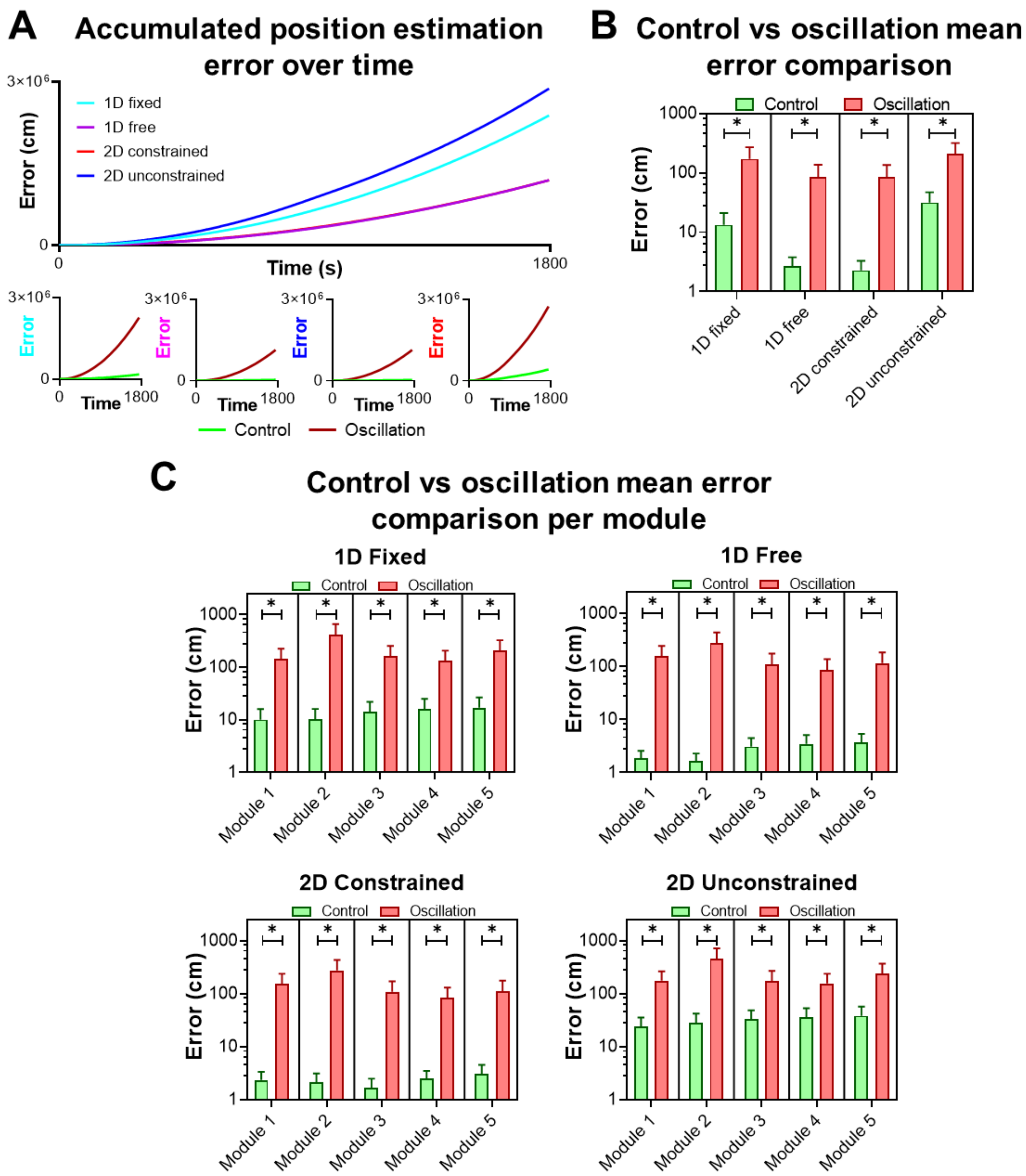


**Supplemental Figure S6. A.** Estimated error accumulation across different experimental conditions. The 1D free and 2D constrained curves are nearly identical and overlap. The four plots below compare trajectory estimation errors with oscillations (red) and without oscillations (green) for each experiment. **B.** Mean estimation error for control and oscillation conditions across all experiments. The left *y*-axis is a $Log_{10}$ scale. Significant differences were found in all cases (Tukey's multiple test, $p < 0.001$). **C.** Mean estimation error comparison between control and oscillation networks, separated by module. The left *y*-axis is a $Log_{10}$ scale. Significant differences were found in all the experimental conditions (Tukey's multiple test, $p < 0.001$).

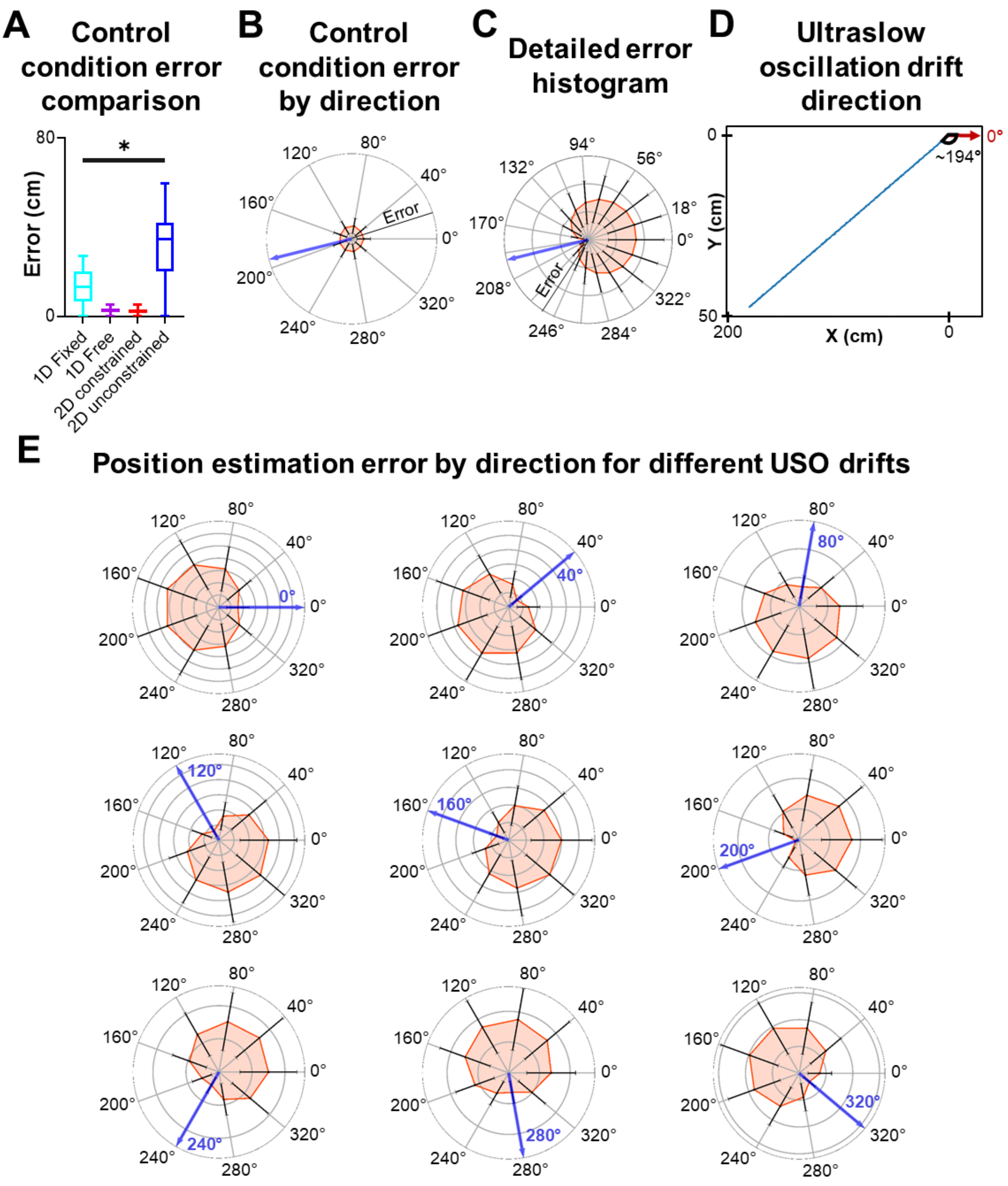


**Supplemental Figure S7. A.** Mean estimation error comparison across experiments under control conditions. Significant differences were observed (One-way ANOVA, $p < 0.001$). **B.** Polar plot showing the mean estimation error of the control network separated by trajectory directions. The radius indicates position estimation error in cm and is divided into sections of 100 cm delimited by grey rings. The blue arrow indicates the drift direction, and the black lines at each point show the standard deviation for each trajectory direction. **C.** Polar plot showing the mean estimation error for different trajectory directions, similar to **Fig. 3F** but using 20 different directions instead of only 10. **D.** Trajectory integrated from the ultraslow oscillation drift. The angle formed by the drift trajectory (blue) with respect to the x-axis (red arrow) is the same as the direction that minimizes position estimation error.

**E.** Polar plots of the mean estimation error for different trajectory directions, similar to **Fig. 2F** but changing oscillation parameters to alter the drift direction caused by the ultraslow oscillation. The blue arrow indicates the drift direction and the radius indicates the position estimation error.

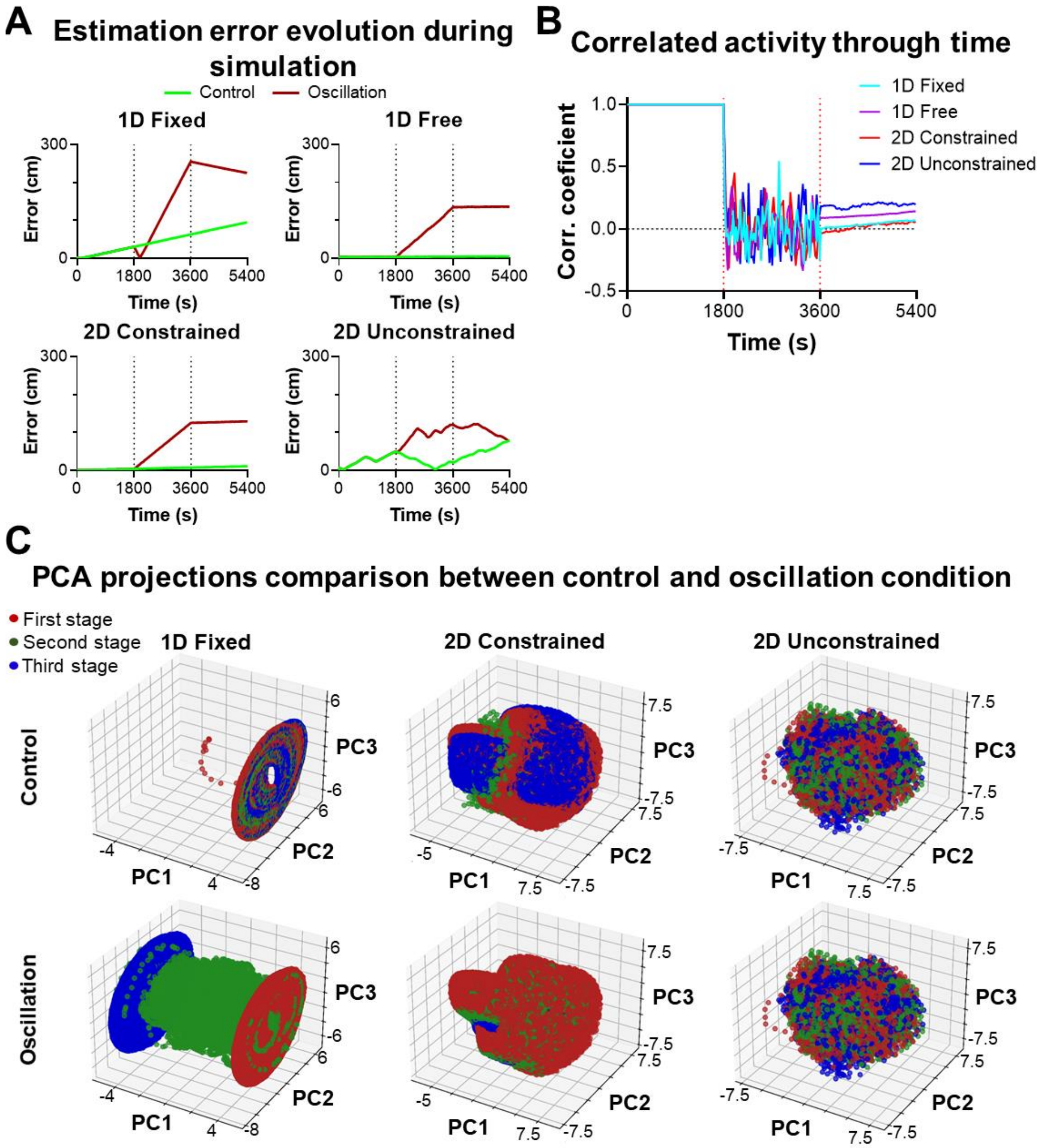


**Supplemental Figure S8. A.** Position estimation error comparison between control (green) and oscillation (red) conditions for each experiment. The three experimental stages are marked with dotted black lines. **B.** Pearson correlation product of control and oscillation activity for each time step of the simulation. The dotted red lines indicate the experimental stages change. **C.** Comparison of the 3D projection of grid cell activity in the PCA space of control and oscillation activity, similar to **Fig. 3A,** but for the remaining experimental conditions. The PCA space where the activity is projected is the same for both conditions. Experimental stages are separated by colors: Start (no oscillation, red); middle (oscillation, green); and end (no oscillation, blue).

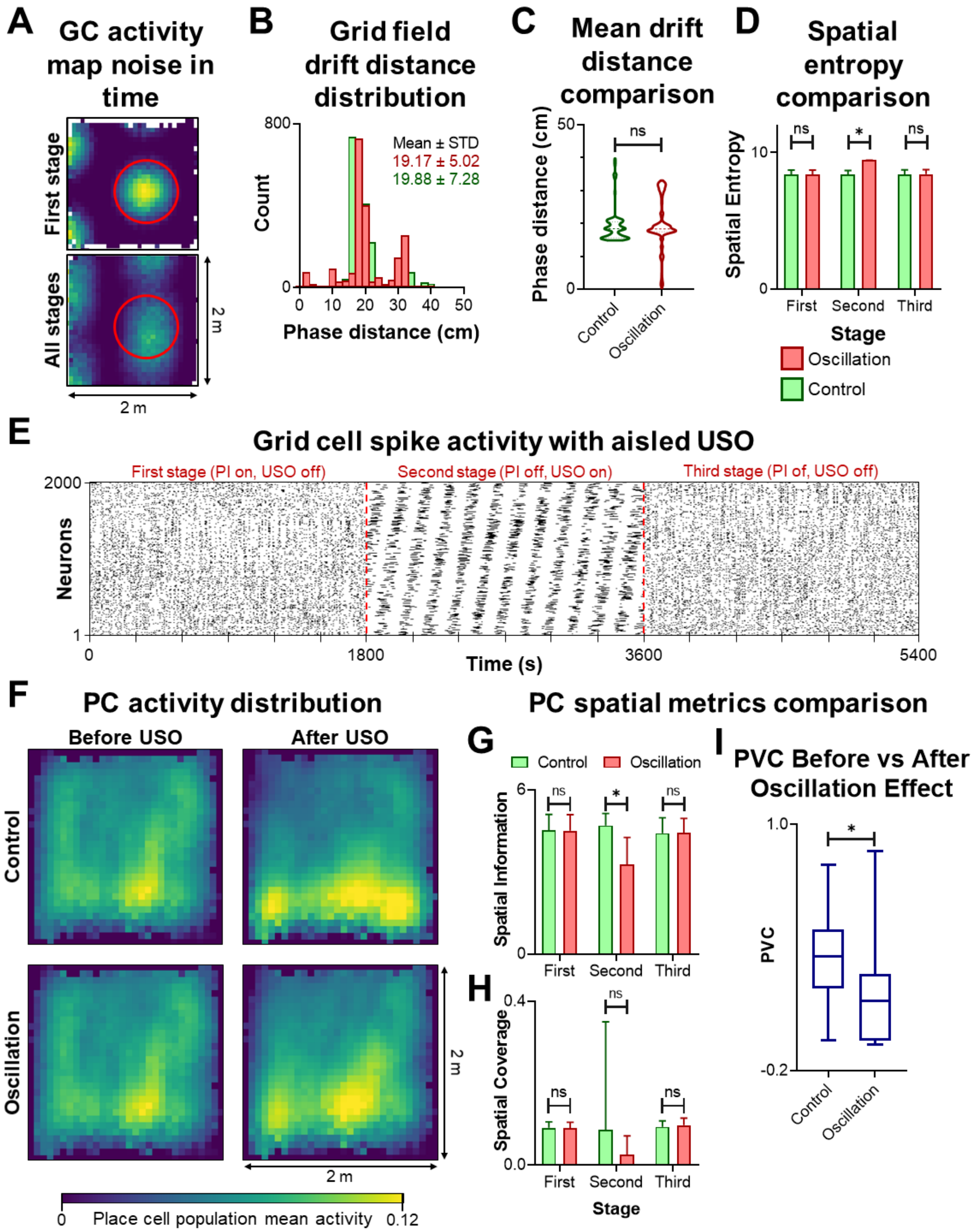


**Supplemental Figure S9. A.** Comparison of grid cell activity map using the recorded activity of only one stage vs using the recordings of the whole experiment. When all experiment stages are used to calculate the activity map, the grid fields enlarge (red circle for size comparison) and become noisier due to the intrinsic path integration error drift. **B.** Histogram of phase distances for control and oscillation conditions, showing the increased standard deviation. **C.** Violin plot comparing grid cells phase distances before and after the effect of ultraslow oscillation in the 2D constrained condition. The mean phase distance of each population shows no significant differences (Wilcoxon test, $p > 0.05$).

**D.** Grid cell spatial entropy comparison between control and oscillatory grid cell networks in the 2D constrained condition, separated by stage. Significant differences were found during the second stage (middle) of the experiment (Dunn's multiple test, $p < 0.001$). For the first (start) and third (end) stage, no significant differences were found (Dunn's multiple test, $p > 0.001$). **E.** Example of grid cell activity dynamics with isolated ultraslow oscillation during the second stage of the experiment. Note that oscillations only appear during the second stage, while in previous experiments (**Supplemental Fig. S5**), oscillations were noticeable during the first and third stages of the experiment. **F.** Mean place cell population activity over the arena, comparing the place field distribution before and after the effects of ultraslow oscillation. **G.** Place cells spatial information and **H.** effective spatial coverage (ESC) comparison between control and oscillation conditions, divided into three experimental stages. No significant differences were found between control and oscillation in each separate stage of the experiment (Dunn's multiple test, $p > 0.05$), except for the spatial information during the second stage of the experiment (Dunn's multiple test, $p < 0.001$). **I.** Population vector correlation (PVC) between place fields before and after the effect of slow oscillations. The PVC was significantly lower in the oscillation condition (Wilcoxon test, $p < 0.001$), indicating a major variation of place fields when compared with the control condition.

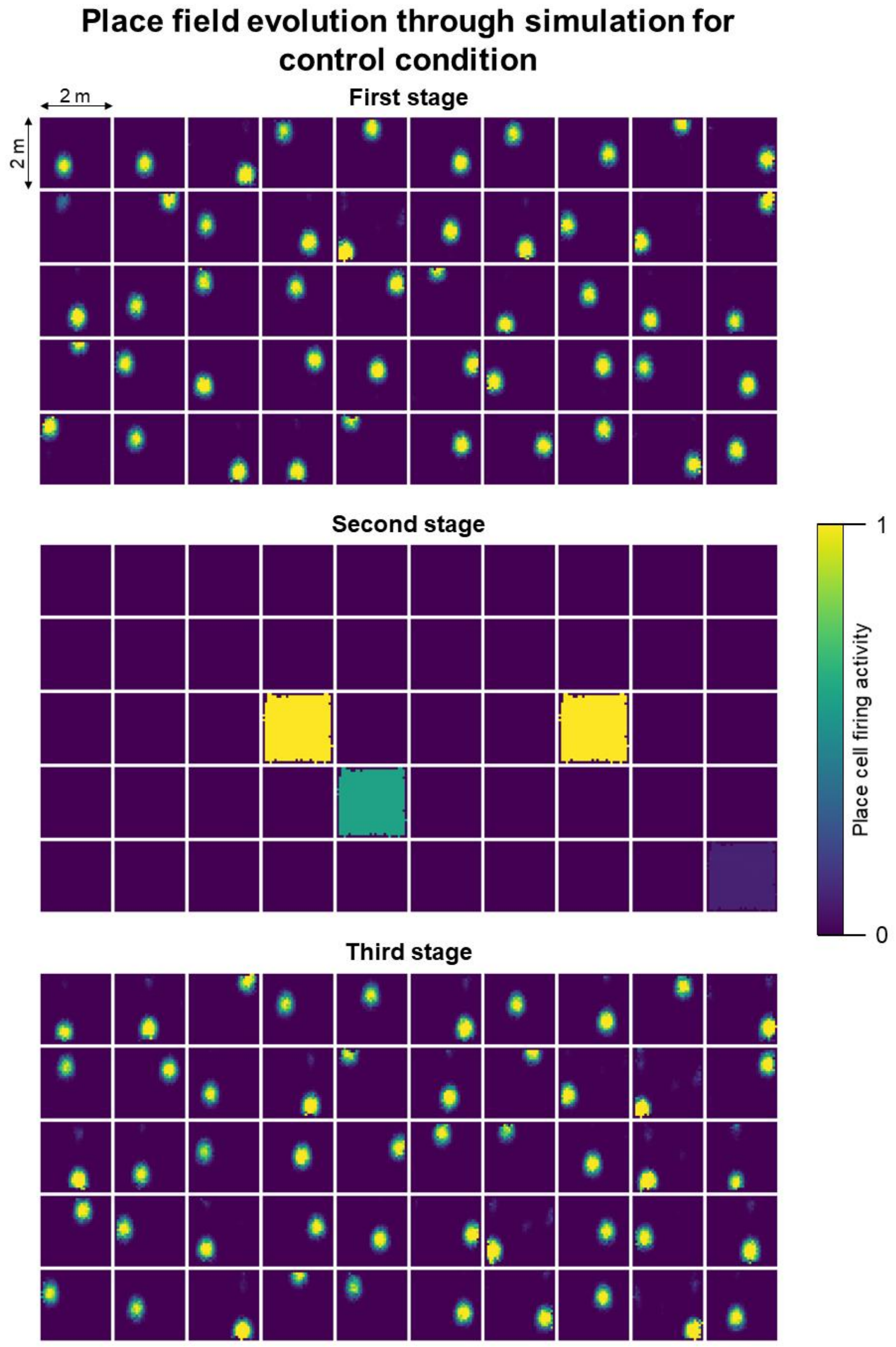


**Supplemental Figure S10.** Place field behavior during each stage of the experiment for the control condition. Each square represents a different place cell. The neuron location in the plot grid is maintained in each stage, meaning that place cell 1 is always plotted in row 1, column 1, place cell 2 in row 1, column 2, and so on. During the second stage, some neurons exhibited a uniform activity map with a firing rate greater than 0. This phenomenon was caused by the state of the network before entering the oscillation period. The control network did not present an ultraslow oscillation; thus, the neurons that were active just prior to the beginning of the second stage remained active thought the whole stage. This phenomenon created the effect that neurons fired through the whole arena when, in reality, it was just path integration being deactivated.

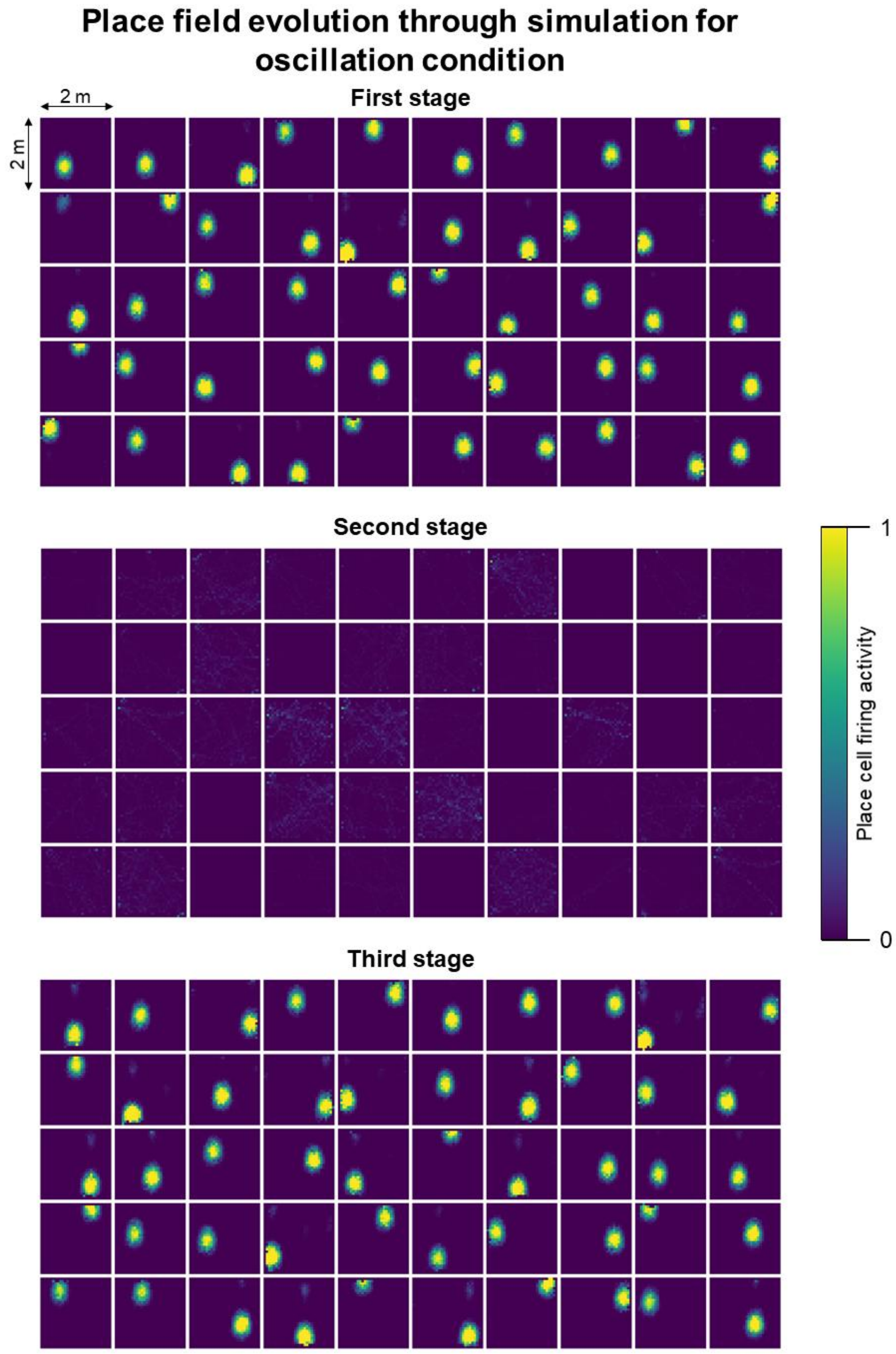


**Supplemental Figure S11.** Place field behavior during each stage of the experiment for the oscillation condition. Each square represents a different place cell. The neuron location in the plot grid is maintained in each stage, meaning that place cell 1 is always plotted in row 1 and column 1, place cell 2 in row 1, column 2, and so on.

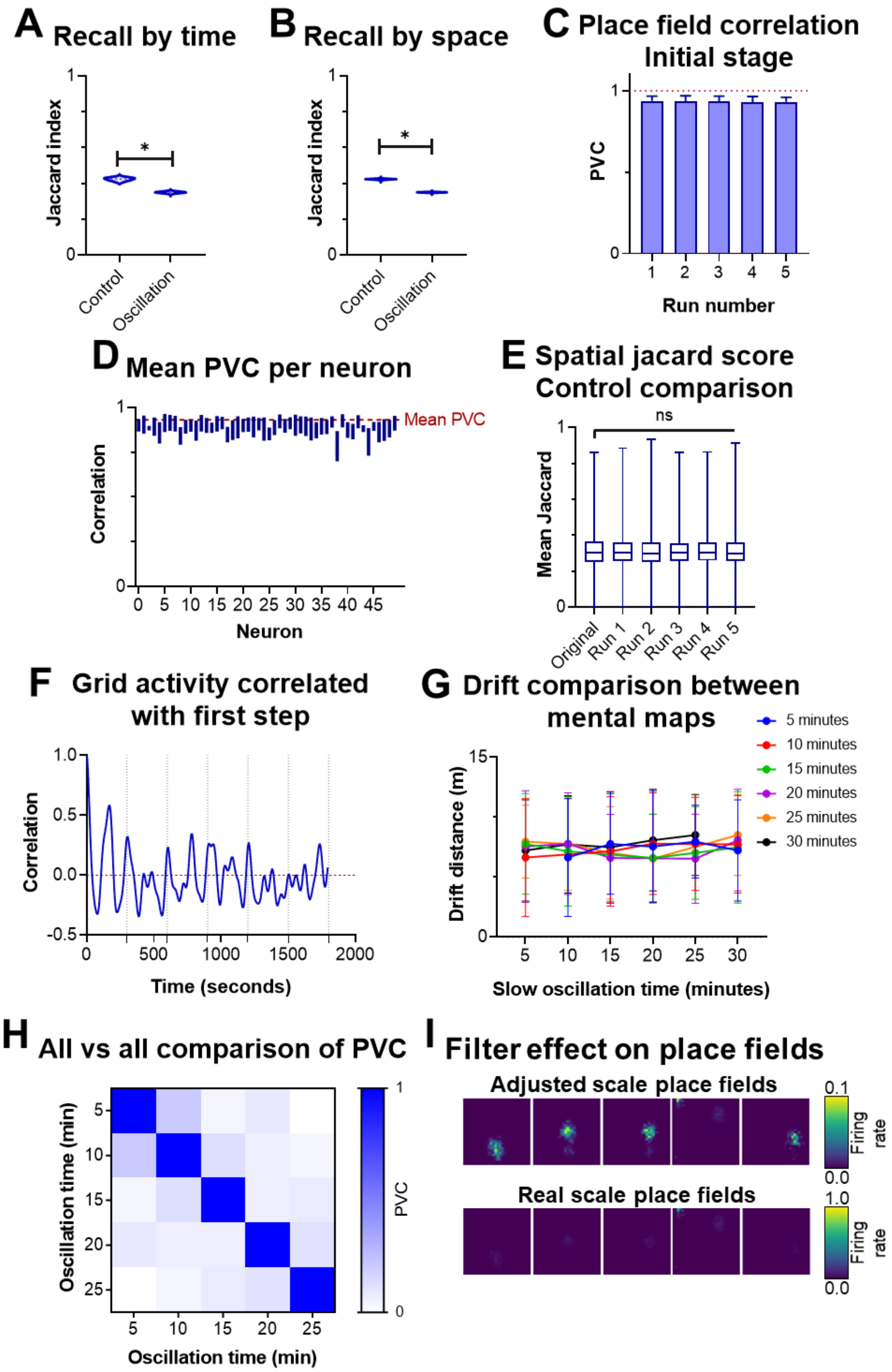


**Supplemental Figure S12. A.** Jaccard similarity index calculated over time and **B.** over location. Significant differences were found between the recalled memory ensemble before and after the ultraslow oscillation period in the control and oscillation conditions (Wilcoxon test, $p < 0.001$). **C.** Population vector correlation (PVC) between the initial conditions of the five new simulations and the original simulation. Red line indicates mean PVC across all neurons. **D.** Mean PVC per place cell across all new simulations. **E.** Jaccard similarity score of the recalled ensembles after the ultraslow oscillation between the new simulations and the original one. A control test was done to assess the Jaccard score without the presence of ultraslow oscillation. In all five simulations, the Jaccard score was low but equal to the control condition (Dunn's multiple comparisons test, $p > 0.05$), indicating no major difference in the recalled memories of the oscillatory condition. **F.** Pearson correlation coefficient between the initial grid cell configuration and the grid cell activity across time. **G.** Drift comparison between the learned place fields of each simulation with different ultraslow oscillation time durations. Differences between the combined time and distance factors were found (Two-way ANOVA, $p < 0.01$) **H.** All-to-all correlation of the learned place fields among all simulations, each with a different ultraslow oscillation duration. **I.** Place fields after the reversed oscillation using the synaptic weights learned after the ultraslow oscillation effect. The scale was adjusted to demonstrate the low place cell activity. When the scale is adjusted to match the other activity maps, place fields disappeared, demonstrating that the reverse oscillation produced a decoupling of the resulting map after the first oscillation period.

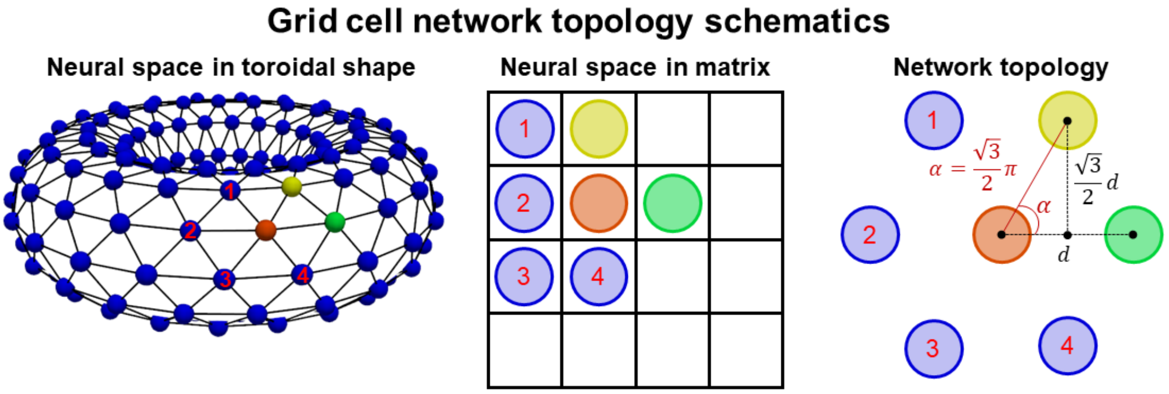


**Supplemental Figure S13.** Schematics of the grid cell network topology. (Left) Grid cells were arranged in a twisted torus, which created the hexagonal shape. (Middle) Computational representation of the grid cell organized in a matrix. Neurons are either numbered or color-coded to show the transformation from the toroidal space to the matrix. (Right) Distances between grid cells in the neural space. Grid cells from the same matrix row were spaced by a distance $d$, while neurons from the same column were spaced by a $\sqrt{3}/2 \ * d$. Grid cells in a row with an even index were shifted by 0.5*$d$ to create an angle of 60° ($\sqrt{3}/2$ * π) between neurons from the same column. This organization created the hexagonal lattice observed in the grid cell activity maps.

**Supplemental Table – Statistical analysis**

**Figure 2C**
**Complete order**

**Friedman test – 1D Fixed, 1D Free, 2D Constrained, 2D Unconstrained**
**P value:** < 0.001
**Are means signif. different? (p < 0.001):** Yes
**Number of groups:** 3
**Friedman statistic:** 60.00, 56.33, 59.36, 60 (Ordered by experimental condition)
**Number of treatments (columns):** 3
**Number of subjects (rows):** 30

**Figure 2C**
**Complete order**

**Dunn's multiple comparisons test – 1D Fixed**
**α = 0.001, R=Rank**

| Stage comparison | R1 | R2 | Rdiff | N1 | N2 | p | p<α? |
|---|---|---|---|---|---|---|---|
| **1 vs 2** | 75.00 | 30.00 | 45.00 | 30 | 30 | <0.0001 | Yes |
| **1 vs 3** | 75.00 | 75.00 | 0.000 | 30 | 30 | >0.9999 | No |
| **2 vs 3** | 30.00 | 75.00 | -45.00 | 30 | 30 | <0.0001 | Yes |

**Figure 2C**
**Complete order**

**Dunn's multiple comparisons test – 1D Free**
**α = 0.001, R=Rank**

| Stage comparison | R1 | R2 | Rdiff | N1 | N2 | p | p<α? |
|---|---|---|---|---|---|---|---|
| **1 vs 2** | 76.00 | 30.00 | 46.00 | 30 | 30 | <0.0001 | Yes |
| **1 vs 3** | 76.00 | 74.00 | 2.000 | 30 | 30 | >0.9999 | No |
| **2 vs 3** | 30.00 | 74.00 | -44.00 | 30 | 30 | <0.0001 | Yes |

**Figure 2C**
**Complete order**

**Dunn's multiple comparisons test – 2D Constrained**
**α = 0.001, R=Rank**

| Stage comparison | R1 | R2 | Rdiff | N1 | N2 | p | p<α? |
|---|---|---|---|---|---|---|---|
| **1 vs 2** | 75.50 | 30.00 | 45.50 | 30 | 30 | <0.0001 | Yes |
| **1 vs 3** | 75.50 | 74.50 | 1.000 | 30 | 30 | >0.9999 | No |
| **2 vs 3** | 30.00 | 74.50 | -44.50 | 30 | 30 | <0.0001 | Yes |

**Figure 2C**
**Complete order**

**Dunn's multiple comparisons test – 2D Unconstrained**
**α = 0.001, R=Rank**

| Stage comparison | R1 | R2 | Rdiff | N1 | N2 | p | p<α? |
|---|---|---|---|---|---|---|---|
| **1 vs 2** | 75.00 | 30.00 | 45.00 | 30 | 30 | <0.0001 | Yes |
| **1 vs 3** | 75.00 | 75.00 | 0.000 | 30 | 30 | >0.9999 | No |
| **2 vs 3** | 30.00 | 75.00 | -45.00 | 30 | 30 | <0.0001 | Yes |

**Figure 2C**
**Immobility order**

**Friedman test – 1D Fixed, 1D Free, 2D Constrained, 2D Unconstrained**
**P value:** < 0.001
**Are means signif. different? (p < 0.001):** Yes
**Number of groups:** 3
**Friedman statistic:** 48.67, 52.77, 58.21, 57.83 (Ordered by experimental condition)
**Number of treatments (columns):** 3
**Number of subjects (rows):** 30

**Figure 2C**
**Immobility order**

**Dunn's multiple comparisons test – 1D Fixed**
**α = 0.001, R=Rank**

| Stage comparison | R1 | R2 | Rdiff | N1 | N2 | p | p<α? |
|---|---|---|---|---|---|---|---|
| **1 vs 2** | 89.00 | 36.50 | 52.50 | 30 | 30 | <0.0001 | Yes |
| **1 vs 3** | 89.00 | 54.50 | 34.50 | 30 | 30 | <0.0001 | Yes |
| **2 vs 3** | 36.50 | 54.50 | -18.00 | 30 | 30 | 0.0604 | No |

**Figure 2C**
**Immobility order**

**Dunn's multiple comparisons test – 1D Free**
**α = 0.001, R=Rank**

| Stage comparison | R1 | R2 | Rdiff | N1 | N2 | p | p<α? |
|---|---|---|---|---|---|---|---|
| **1 vs 2** | 88.50 | 33.00 | 55.50 | 30 | 30 | <0.0001 | Yes |
| **1 vs 3** | 88.50 | 58.50 | 30.00 | 30 | 30 | 0.0003 | Yes |
| **2 vs 3** | 33.00 | 58.50 | -25.50 | 30 | 30 | 0.0030 | No |

**Figure 2C**
**Immobility order**

**Dunn's multiple comparisons test – 2D Constrained**
**α = 0.001, R=Rank**

| Stage comparison | R1 | R2 | Rdiff | N1 | N2 | p | p<α? |
|---|---|---|---|---|---|---|---|
| **1 vs 2** | 88.00 | 30.00 | 58.00 | 30 | 30 | <0.0001 | Yes |
| **1 vs 3** | 88.00 | 62.00 | 26.00 | 30 | 30 | 0.0024 | No |
| **2 vs 3** | 30.00 | 62.00 | -32.00 | 30 | 30 | 0.0001 | Yes |

**Figure 2C**
**Immobility order**

**Dunn's multiple comparisons test – 2D Unconstrained**
**α = 0.001, R=Rank**

| Stage comparison | R1 | R2 | Rdiff | N1 | N2 | p | p<α? |
|---|---|---|---|---|---|---|---|
| **1 vs 2** | 87.50 | 30.00 | 57.50 | 30 | 30 | <0.0001 | Yes |
| **1 vs 3** | 87.50 | 62.50 | 25.00 | 30 | 30 | 0.0037 | No |
| **2 vs 3** | 30.00 | 62.50 | -32.50 | 30 | 30 | <0.0001 | Yes |

**Figure 2C**
**Movement order**

**Friedman test – 1D Fixed, 1D Free, 2D Constrained, 2D Unconstrained**
**P value:** < 0.001
**Are means signif. different? (p < 0.001)** Yes
**Number of groups:** 3
**Friedman statistic:** 22.84, 37.38, 48.49, 46.85 (Ordered by experimental condition)
**Number of treatments (columns):** 3
**Number of subjects (rows):** 30

**Figure 2C**
**Movement order**

**Dunn's multiple comparisons test – 1D Fixed**
**α = 0.001, R=Rank**

| Stage comparison | R1 | R2 | Rdiff | N1 | N2 | p | p<α? |
|---|---|---|---|---|---|---|---|
| **1 vs 2** | 67.50 | 39.00 | 28.50 | 30 | 30 | 0.0007 | Yes |
| **1 vs 3** | 67.50 | 73.50 | -6.000 | 30 | 30 | >0.9999 | No |
| **2 vs 3** | 39.00 | 73.50 | -34.50 | 30 | 30 | <0.0001 | Yes |

**Figure 2C**
**Movement order**

**Dunn's multiple comparisons test – 1D Free**
**α = 0.001, R=Rank**

| Stage comparison | R1 | R2 | Rdiff | N1 | N2 | p | p<α? |
|---|---|---|---|---|---|---|---|
| **1 vs 2** | 73.50 | 33.00 | 40.50 | 30 | 30 | <0.0001 | Yes |
| **1 vs 3** | 73.50 | 73.50 | 0.000 | 30 | 30 | >0.9999 | No |
| **2 vs 3** | 33.00 | 73.50 | -40.50 | 30 | 30 | <0.0001 | Yes |

**Figure 2C**
**Movement order**

**Dunn's multiple comparisons test – 2D Constrained**
**α = 0.001, R=Rank**

| Stage comparison | R1 | R2 | Rdiff | N1 | N2 | p | p<α? |
|---|---|---|---|---|---|---|---|
| **1 vs 2** | 79.00 | 30.00 | 49.00 | 30 | 30 | <0.0001 | Yes |
| **1 vs 3** | 79.00 | 71.00 | 8.000 | 30 | 30 | 0.9051 | No |
| **2 vs 3** | 30.00 | 71.00 | -41.00 | 30 | 30 | <0.0001 | Yes |

**Figure 2C**
**Movement order**

**Dunn's multiple comparisons test – 2D Unconstrained**
**α = 0.001, R=Rank**

| Stage comparison | R1 | R2 | Rdiff | N1 | N2 | p | p<α? |
|---|---|---|---|---|---|---|---|
| **1 vs 2** | 79.00 | 30.00 | 49.00 | 30 | 30 | <0.0001 | Yes |
| **1 vs 3** | 79.00 | 71.00 | 8.000 | 30 | 30 | 0.9051 | No |
| **2 vs 3** | 30.00 | 71.00 | -41.00 | 30 | 30 | <0.0001 | Yes |

**Figure 2E**

**ANOVA summary**
**F:** 7773
**P value:** <0.0001
**Significant diff. among means (P < 0.001)?** Yes
**R square:** 0.2953
**Number of treatments (columns):** 4
**Number of values (total):** 55656

**Figure 2E**

**Tukey's multiple comparisons test**
**α = 0.001**

| Stage comparison | Mean Diff. | SE of diff. | N1-2 | p | p<α? |
|---|---|---|---|---|---|
| **1D fixed vs. 1D free** | 85.83 | 0.9881 | 13914 | <0.0001 | Yes |
| **1D fixed vs. 2D constr.** | 85.83 | 0.9881 | 13914 | <0.0001 | Yes |
| **1D fixed vs. 2D uncon.** | -35.68 | 0.9881 | 13914 | <0.0001 | Yes |
| **1D free vs. 2D constr.** | -0.007203 | 0.9881 | 13914 | >0.9999 | No |
| **1D free vs. 2D uncon.** | -121.5 | 0.9881 | 13914 | <0.0001 | Yes |
| **2D constr. vs. 2D uncon.** | -121.5 | 0.9881 | 13914 | <0.0001 | Yes |

**Figure 2F**

**One-way ANOVA – Comparison between direction angles**
**F:** 8804
**P value:** <0.0001
**Significant diff. among means (P < 0.001)?** Yes
**R square:** 0.3600
**Number of treatments (columns):** 9
**Number of values (total):** 125226

**Figure 2F**

**Dunnett's multiple comparison test - 200° vs all angle**
**α = 0.001**

| Angle comparison | Mean Diff. | SE of diff. | N1-2 | p | p<α? |
|---|---|---|---|---|---|
| **200 vs. 0** | -162.0 | 0.8514 | 13914 | <0.0001 | Yes |
| **200 vs. 40** | -157.9 | 0.8514 | 13914 | <0.0001 | Yes |
| **200 vs. 80** | -135.7 | 0.8514 | 13914 | <0.0001 | Yes |
| **200 vs. 120** | -95.70 | 0.8514 | 13914 | <0.0001 | Yes |
| **200 vs. 160** | -43.06 | 0.8514 | 13914 | <0.0001 | Yes |
| **200 vs. 240** | -56.68 | 0.8514 | 13914 | <0.0001 | Yes |
| **200 vs. 280** | -107.4 | 0.8514 | 13914 | <0.0001 | Yes |
| **200 vs. 320** | -143.2 | 0.8514 | 13914 | <0.0001 | Yes |

**Figure 3C**

**Dunn's multiple comparisons test – Control vs Oscillation**
**α = 0.001, R=Rank**

| Stage | R1 | R2 | Rdiff | N1 | N2 | p | p<α? |
|---|---|---|---|---|---|---|---|
| **1** | 7843 | 7843 | 0.000 | 2000 | 2000 | >0.9999 | No |
| **2** | 8983 | 2000 | 6983 | 2000 | 2000 | <0.0001 | Yes |
| **3** | 7823 | 7508 | 315.0 | 2000 | 2000 | 0.0233 | No |

**Figure 3D**

**Dunn's multiple comparisons test – Control vs Oscillation**
**α = 0.001, R=Rank**

| Stage | R1 | R2 | Rdiff | N1 | N2 | p | p<α? |
|---|---|---|---|---|---|---|---|
| **1** | 5897 | 5897 | 0.000 | 2000 | 2000 | >0.9999 | No |
| **2** | 4832 | 12000 | -7169 | 2000 | 2000 | <0.0001 | Yes |
| **3** | 6560 | 6815 | -255.5 | 2000 | 2000 | 0.0925 | No |

**Figure 3F**

**Dunn's multiple comparisons - Oscillation condition**
**α = 0.001, R=Rank**

| Stage comparison | R1 | R2 | Rdiff | N1 | N2 | p | p<α? |
|---|---|---|---|---|---|---|---|
| **1 vs 2** | 127.0 | 50.00 | 77.00 | 50 | 50 | <0.0001 | Yes |
| **1 vs 3** | 127.0 | 123.0 | 4.000 | 50 | 50 | >0.9999 | No |
| **2 vs 3** | 50.00 | 123.0 | -73.00 | 50 | 50 | <0.0001 | Yes |

**Figure 3G**

**Wilcoxon matched-pairs test – First vs Third stage**
**P value:** 0.0020
**Significantly different (p < 0.001)?** No
**Significantly different (p < 0.01)?** Yes
**One- or two-tailed p value?** Two-tailed
**Sum of positive, negative ranks:** 883.0, -293.0
**Sum of signed ranks (W):** 590.0
**Number of pairs:** 50
**Number of ties (ignored):** 2

**Figure 4A**

**Wilcoxon matched-pairs test – Oscillation vs Control**
**P value:** 0.0001
**Significantly different (P < 0.001)?** Yes
**One- or two-tailed P value?** Two-tailed
**Sum of positive, negative ranks:** 1019, -256.0
**Sum of signed ranks (W):** 763.0
**Number of pairs:** 50
**Number of ties (ignored):** 0

**Figure 4A**

**Pitman-Morgan test – Oscillation vs Control**
**P value:** 0.0917
**Significantly different (P < 0.001)?** No
**One- or two-tailed P value?** Two-tailed
**Degrees of freedom:** 48
**Sample size:** 50

**Figure 4B**

**Pitman-Morgan test – Oscillation vs Control**
**P value:** <0.0001
**Significantly different (P < 0.001)?** Yes
**One- or two-tailed P value?** Two-tailed
**Degrees of freedom:** 48
**Sample size:** 50

**Figure 4E**

**Friedman test – Duration comparison**
**P value:** 0.1450
**Are means signif. different? (P < 0.05)** No
**Number of groups:** 5
**Friedman statistic:** 6.832
**Number of treatments (columns):** 5
**Number of subjects (rows):** 50

**Figure 4E**

**One sample t-test**
**Theoretical mean = 0, α = 0.05**

| Osc. duration | Mean | t | df | n | p | p<α |
|---|---|---|---|---|---|---|
| **5** | 0.042 | 1.50 | 49 | 50 | 0.139 | No |
| **10** | 0.034 | 1.07 | 49 | 50 | 0.287 | No |
| **15** | 0.088 | 2.41 | 49 | 50 | 0.0196 | Yes |
| **20** | 0.037 | 1.19 | 49 | 50 | 0.2398 | No |
| **25** | -0.018 | 1 | 49 | 50 | 0.3220 | No |

**Figure 4F**

**Friedman test – Synaptic weight comparison**
**P value:** <0.0001
**Are means signif. different? (P < 0.001):** Yes
**Number of groups:** 3
**Friedman statistic:** 75.64
**Number of treatments (columns):** 3
**Number of subjects (rows):** 50

**Figure 4F**

**Dunn's multiple comparisons - Oscillation condition**
**α = 0.001, β = 0.05, R = Rank**

| W comparison | R1 | R2 | Rdiff | N1 | N2 | p | p<α? | p>β? |
|---|---|---|---|---|---|---|---|---|
| W1 vs. W2 | 79.00 | 150.0 | -71.00 | 50 | 50 | <0.0001 | Yes | No |
| W1 vs. W0 | 79.00 | 71.00 | 8.000 | 50 | 50 | >0.9999 | No | Yes |
| W2 vs. W0 | 150.0 | 71.00 | 79.00 | 50 | 50 | <0.0001 | Yes | No |

**Supplemental Figure S3C**

**Wilcoxon matched-pairs test – First vs second stage**
**P value:** <0.0001
**Significantly different (P < 0.001)?** Yes
**One- or two-tailed P value?** Two-tailed
**Sum of positive, negative ranks:** 0.000, -210.0
**Sum of signed ranks (W):** -210.0
**Number of pairs:** 20
**Number of ties (ignored):** 0

**Supplemental Figure S3D 1D Fixed**

**Friedman test**
**P value:** < 0.0001
**Are means signif. different? (p < 0.001):** Yes
**Number of groups:** 3
**Friedman statistic:** 57
**Number of treatments (columns):** 3
**Number of subjects (rows):** 30

**Supplemental Figure S3D 1D Fixed**

**Dunn's multiple comparisons test – 1D Fixed**
**α = 0.001, R=Rank**

| Order comparison | R1 | R2 | Rdiff | N1 | N2 | p | p<α? |
|---|---|---|---|---|---|---|---|
| **Mov. vs Still** | 30.00 | 78.00 | -48.00 | 30 | 30 | <0.0001 | Yes |
| **Mov. vs Total** | 30.00 | 72.00 | -42.00 | 30 | 30 | <0.0001 | Yes |
| **Still vs Total** | 78.00 | 72.00 | 6.000 | 30 | 30 | >0.9999 | No |

**Supplemental Figure S3D 1D Free**

**Friedman test**
**P value:** < 0.0001
**Are means signif. different? (p < 0.001):** Yes
**Number of groups:** 3
**Friedman statistic:** 56.48
**Number of treatments (columns):** 3
**Number of subjects (rows):** 30

**Supplemental Figure S3D 1D Free**

**Dunn's multiple comparisons test – 1D Fixed**
**α = 0.001, R=Rank**

| Order comparison | R1 | R2 | Rdiff | N1 | N2 | p | p<α? |
|---|---|---|---|---|---|---|---|
| **Mov. vs Still** | 30.00 | 87.00 | -57.00 | 30 | 30 | <0.0001 | Yes |
| **Mov. vs Total** | 30.00 | 63.00 | -33.00 | 30 | 30 | <0.0001 | Yes |
| **Still vs Total** | 87.00 | 63.00 | 24.00 | 30 | 30 | 0.0058 | No |

**Supplemental Figure S3D 2D Constrained**

**Friedman test**
**P value:** < 0.0001
**Are means signif. different? (p < 0.001):** Yes
**Number of groups:** 3
**Friedman statistic:** 54.60
**Number of treatments (columns):** 3
**Number of subjects (rows):** 30

**Supplemental Figure S3D 2D Constrained**

**Dunn's multiple comparisons test – 1D Fixed**
**α = 0.001, R=Rank**

| Order comparison | R1 | R2 | Rdiff | N1 | N2 | p | p<α? |
|---|---|---|---|---|---|---|---|
| **Mov. vs Still** | 30.00 | 87.00 | -57.00 | 30 | 30 | <0.0001 | Yes |
| **Mov. vs Total** | 30.00 | 63.00 | -33.00 | 30 | 30 | <0.0001 | Yes |
| **Still vs Total** | 87.00 | 63.00 | 24.00 | 30 | 30 | 0.0058 | No |

**Supplemental Figure S3D 2D Unconstrained**

**Friedman test**
**P value:** < 0.0001
**Are means signif. different? (p < 0.001):** Yes
**Number of groups:** 3
**Friedman statistic:** 47.95
**Number of treatments (columns):** 3
**Number of subjects (rows):** 30

**Supplemental Figure S3D 2D Unconstrained**

**Dunn's multiple comparisons test – 1D Fixed**
**α = 0.001, R=Rank**

| Order comparison | R1 | R2 | Rdiff | N1 | N2 | p | p<α? |
|---|---|---|---|---|---|---|---|
| **Mov. vs Still** | 30.00 | 76.50 | -46.50 | 30 | 30 | <0.0001 | Yes |
| **Mov. vs Total** | 30.00 | 73.50 | -43.50 | 30 | 30 | <0.0001 | Yes |
| **Still vs Total** | 76.50 | 73.50 | 3.000 | 30 | 30 | >0.9999 | No |

**Supplemental Figure S6B**

**Repeated measures ANOVA summary**
**Assume sphericity?** No
**F:** 45116
**P value:** <0.0001
**Statistically significant (P < 0.001)?** Yes
**Geisser-Greenhouse's epsilon:** 0.1445
**R square:** 0.7643
**Number of treatments (columns):** 8
**Number of subjects (rows):** 13914
**Number of missing values:** 0

**Supplemental Figure S6B**

**Tukey's multiple comparisons test – Control vs Oscillation**
**α = 0.001**

| Stage comparison | Mean Diff. | SE of diff. | N1-2 | p | p<α? |
|---|---|---|---|---|---|
| **1D Fixed** | -158.9 | 0.7781 | 13914 | <0.0001 | Yes |
| **1D Free** | -83.83 | 0.4243 | 13914 | <0.0001 | Yes |
| **2D Constrained** | -84.20 | 0.4166 | 13914 | <0.0001 | Yes |
| **2D Unconstrained** | -176.3 | 0.8067 | 13914 | <0.0001 | Yes |

**Supplemental Figure S6C 1D Fixed**

**Repeated measures ANOVA summary**
**Assume sphericity?** No
**F:** 41716
**P value:** <0.0001
**Statistically significant (P < 0.001)?** Yes
**Geisser-Greenhouse's epsilon:** 0.1111
**R square:** 0.7499

**Number of treatments (columns):** 10
**Number of subjects (rows):** 13914
**Number of missing values:** 0

**Supplemental Figure S6C 1D Fixed**

**Tukey's multiple comparisons test – Comparison by module**
**α = 0.001**

| Grid module | Mean Diff. | SE of diff. | N1-2 | p | p<α? |
|---|---|---|---|---|---|
| **M1** | -132.7 | 0.6479 | 13914 | <0.0001 | Yes |
| **M2** | -409.8 | 2.007 | 13914 | <0.0001 | Yes |
| **M3** | -146.7 | 0.7192 | 13914 | <0.0001 | Yes |
| **M4** | -115.2 | 0.5645 | 13914 | <0.0001 | Yes |
| **M5** | -189.0 | 0.9261 | 13914 | <0.0001 | Yes |

**Supplemental Figure S6C 1D Free**

**Repeated measures ANOVA summary**
**Assume sphericity?** No
**F:** 40647
**P value:** <0.0001
**Statistically significant (P < 0.001)?** Yes
**Geisser-Greenhouse's epsilon:** 0.1111
**R square:** 0.7450
**Number of treatments (columns):** 10
**Number of subjects (rows):** 13914
**Number of missing values:** 0

**Supplemental Figure S6C 1D Free**

**Tukey's multiple comparisons test – Comparison by module**
**α = 0.001**

| Grid module | Mean Diff. | SE of diff. | N1-2 | p | p<α? |
|---|---|---|---|---|---|
| **M1** | -153.8 | 0.7550 | 13914 | <0.0001 | Yes |
| **M2** | -276.7 | 1.374 | 13914 | <0.0001 | Yes |
| **M3** | -107.6 | 0.5371 | 13914 | <0.0001 | Yes |
| **M4** | -83.36 | 0.4162 | 13914 | <0.0001 | Yes |
| **M5** | -111.9 | 0.5631 | 13914 | <0.0001 | Yes |

**Supplemental Figure S6C 2D Constrained**

**Repeated measures ANOVA summary**
**Assume sphericity?** No
**F:** 41238
**P value:** <0.0001
**Statistically significant (P < 0.001)?** Yes
**Geisser-Greenhouse's epsilon:** 0.1111
**R square:** 0.7477
**Number of treatments (columns):** 10
**Number of subjects (rows):** 13914
**Number of missing values:** 0

**Supplemental Figure S6C 2D Constrained**

**Tukey's multiple comparisons test – Comparison by module**
**α = 0.001**

| Grid module | Mean Diff. | SE of diff. | N1-2 | p | p<α? |
|---|---|---|---|---|---|
| **M1** | -148.5 | 0.7264 | 13914 | <0.0001 | Yes |
| **M2** | -271.1 | 1.337 | 13914 | <0.0001 | Yes |
| **M3** | -106.8 | 0.5281 | 13914 | <0.0001 | Yes |
| **M4** | -79.88 | 0.3950 | 13914 | <0.0001 | Yes |
| **M5** | -108.5 | 0.5323 | 13914 | <0.0001 | Yes |

<table>
<tr><td>Supplemental<br>Figure S6C<br>2D Unconstrained</td><td>Repeated measures ANOVA summary<br>Assume sphericity? No<br>F: 43157<br>P value: <0.0001<br>Statistically significant (P < 0.001)? Yes<br>Geisser-Greenhouse's epsilon: 0.1115<br>R square: 0.7562<br>Number of treatments (columns): 10<br>Number of subjects (rows): 13914<br>Number of missing values: 0</td></tr>
<tr><td>Supplemental<br>Figure S6C<br>2D Unconstrained</td><td>Tukey's multiple comparisons test – Comparison by module<br>α = 0.001
<table>
<tr><th>Grid module</th><th>Mean Diff.</th><th>SE of diff.</th><th>N1-2</th><th>p</th><th>p<α?</th></tr>
<tr><td>M1</td><td>-151.0</td><td>0.6481</td><td>13914</td><td><0.0001</td><td>Yes</td></tr>
<tr><td>M2</td><td>-425.3</td><td>2.068</td><td>13914</td><td><0.0001</td><td>Yes</td></tr>
<tr><td>M3</td><td>-140.1</td><td>0.6780</td><td>13914</td><td><0.0001</td><td>Yes</td></tr>
<tr><td>M4</td><td>-119.9</td><td>0.5410</td><td>13914</td><td><0.0001</td><td>Yes</td></tr>
<tr><td>M5</td><td>-201.9</td><td>0.9242</td><td>13914</td><td><0.0001</td><td>Yes</td></tr>
</table></td></tr>
<tr><td>Supplemental<br>Figure S7A</td><td>ANOVA summary<br>F: 35689<br>P value: <0.0001<br>Significant diff. among means (P < 0.001)? Yes<br>R square: 0.6580<br>Number of treatments (columns): 4<br>Number of values (total): 55656</td></tr>
<tr><td>Supplemental<br>Figure S7A</td><td>Tukey's multiple comparisons test<br>α = 0.001
<table>
<tr><th>Stage comparison</th><th>Mean Diff.</th><th>SE of diff.</th><th>N1-2</th><th>p</th><th>p<α?</th></tr>
<tr><td>1D fixed vs. 1D free</td><td>10.72</td><td>0.1034</td><td>13914</td><td><0.0001</td><td>Yes</td></tr>
<tr><td>1D fixed vs. 2D constr.</td><td>11.09</td><td>0.1034</td><td>13914</td><td><0.0001</td><td>Yes</td></tr>
<tr><td>1D fixed vs. 2D uncon.</td><td>-18.36</td><td>0.1034</td><td>13914</td><td><0.0001</td><td>Yes</td></tr>
<tr><td>1D free vs. 2D constr.</td><td>0.3674</td><td>0.1034</td><td>13914</td><td>0.0021</td><td>No</td></tr>
<tr><td>1D free vs. 2D uncon.</td><td>-29.08</td><td>0.1034</td><td>13914</td><td><0.0001</td><td>Yes</td></tr>
<tr><td>2D constr. vs. 2D uncon.</td><td>-29.45</td><td>0.1034</td><td>13914</td><td><0.0001</td><td>Yes</td></tr>
</table></td></tr>
<tr><td>Supplemental<br>Figure S7C</td><td>ANOVA summary<br>F: 8008<br>P value: <0.0001<br>Significant diff. among means (P < 0.001)? Yes<br>R square: 0.3529<br>Number of treatments (columns): 19<br>Number of values (total): 264366</td></tr>
<tr><td>Supplemental<br>Figure S9C</td><td>Wilcoxon matched-pairs signed rank test<br>P value: 0.3306<br>Significantly different (P < 0.05)? No<br>One- or two-tailed P value? Two-tailed<br>Sum of positive, negative ranks: 948650, -901277<br>Sum of signed ranks (W): 47373<br>Number of pairs: 2000<br>Number of ties (ignored): 77</td></tr>
</table>

| | |
|---|---|
| **Supplemental Figure S9C** | **Pitman-Morgan test – Oscillation vs Control**<br>**P value:** <0.0001<br>**Significantly different (P < 0.001)?** Yes<br>**One- or two-tailed P value?** Two-tailed<br>**Degrees of freedom:** 1998<br>**Sample size:** 2000 |
| **Supplemental Figure S9D** | **Dunn's multiple comparisons test – Control vs Oscillation**<br>**α = 0.001, R=Rank** |

| Stage | R1 | R2 | Rdiff | N1 | N2 | p | p<α? |
|---|---|---|---|---|---|---|---|
| **1** | 5944 | 5944 | 0.000 | 2000 | 2000 | >0.9999 | No |
| **2** | 4641 | 12000 | -7359 | 2000 | 2000 | <0.0001 | Yes |
| **3** | 6539 | 6932 | -393.0 | 2000 | 2000 | 0.0027 | No |

| | |
|---|---|
| **Supplemental Figure S9G** | **Dunn's multiple comparisons test – Control vs Oscillation**<br>**α = 0.001, R=Rank** |

| Stage | R1 | R2 | Rdiff | N1 | N2 | p | p<α? |
|---|---|---|---|---|---|---|---|
| **1** | 197.0 | 198.0 | -1.000 | 50 | 50 | >0.9999 | No |
| **2** | 221.0 | 87.00 | 134.0 | 50 | 50 | <0.0001 | Yes |
| **3** | 175.0 | 172.0 | 3.000 | 50 | 50 | >0.9999 | No |

| | |
|---|---|
| **Supplemental Figure S9H** | **Dunn's multiple comparisons test – Control vs Oscillation**<br>**α = 0.001, R=Rank** |

| Stage | R1 | R2 | Rdiff | N1 | N2 | p | p<α? |
|---|---|---|---|---|---|---|---|
| **1** | 199.0 | 200.5 | -1.500 | 50 | 50 | >0.9999 | No |
| **2** | 79.50 | 109.5 | -30.00 | 50 | 50 | 0.3264 | No |
| **3** | 216.0 | 245.5 | -29.50 | 50 | 50 | 0.3445 | No |

| | |
|---|---|
| **Supplemental Figure S9I** | **Wilcoxon matched-pairs signed rank test**<br>**P value:** <0.0001<br>**Significantly different (P < 0.001)?** Yes<br>**One- or two-tailed P value?** Two-tailed<br>**Sum of positive, negative ranks:** 189.0, -1086<br>**Sum of signed ranks (W):** -897.0<br>**Number of pairs:** 50<br>**Number of ties (ignored):** 0 |
| **Supplemental Figure S12A** | **One sample t test - Control**<br>**Theoretical mean:** 1.000<br>**Actual mean:** 0.4236<br>**Number of values:** 13914<br>**T:**7006, **df:**13913<br>**P value (two tailed):** <0.0001<br>**Significant (alpha=0.001)?** Yes |
| **Supplemental Figure S12A** | **Wilcoxon matched-pairs signed rank test**<br>**P value:** <0.0001<br>**Significantly different (P < 0.001)?** Yes<br>**One- or two-tailed P value?** Two-tailed<br>**Sum of positive, negative ranks:** 0.000, -96806655<br>**Sum of signed ranks (W):** -96806655<br>**Number of pairs:** 13914<br>**Number of ties (ignored):** 0 |

**Supplemental Figure S12B**

**Wilcoxon matched-pairs signed rank test**
**P value:** <0.0001
**Significantly different (P < 0.001)?** Yes
**One- or two-tailed P value?** Two-tailed
**Sum of positive, negative ranks:** 0.000, -306153
**Sum of signed ranks (W):** -306153
**Number of pairs:** 782
**Number of ties (ignored):** 0

**Supplemental Figure S12E**

**Dunn's multiple comparisons test – Control vs Run iteration**
**α = 0.001, R=Rank**

| Run | R1 | R2 | Rdiff | N1 | N2 | p | p<α? |
|---|---|---|---|---|---|---|---|
| **1** | 48901 | 49108 | -207.0 | 13914 | 13914 | >0.9999 | No |
| **2** | 48901 | 47962 | 939.0 | 13914 | 13914 | 0.0131 | No |
| **3** | 48901 | 48730 | 171.0 | 13914 | 13914 | >0.9999 | No |
| **4** | 48901 | 49078 | -177.0 | 13914 | 13914 | >0.9999 | No |
| **5** | 48901 | 48418 | 483.0 | 13914 | 13914 | 0.6085 | No |

**Supplemental Figure S12G**

**Two-way ANOVA – Time and drift distance comparison**

| Source of Variation | % of total variation | P value | Significant? |
|---|---|---|---|
| **Interaction** | 36.36 | <0.0001 | Yes |
| **Row Factor** | 0.1163 | 0.6639 | No |
| **Column Factor** | 0.1163 | 0.6639 | No |

| ANOVA table | SS | DF | MS | F (DFn, DFd) | P value |
|---|---|---|---|---|---|
| **Interaction** | 14291 | 25 | 571.7 | F (25, 1764) = 40.47 | P<0.0001 |
| **Row Factor** | 45.70 | 5 | 9.140 | F (5, 1764) = 0.6470 | P=0.6639 |
| **Column Factor** | 45.70 | 5 | 9.140 | F (5, 1764) = 0.6470 | P=0.6639 |
| **Residual** | 24920 | 1764 | 14.13 | | |
